# From Noise to Models to Numbers: Evaluating Negative Binomial Models and Parameter Estimations in Single-Cell RNA-seq

**DOI:** 10.1101/2025.05.05.652189

**Authors:** Yiling Wang, Zhanpeng Shu, Zhixing Cao, Ramon Grima

## Abstract

The Negative Binomial (NB) distribution is widely used to approximate transcript count distributions in single-cell RNA sequencing (scRNA-seq) data, yet the reason for its ubiquity is not fully understood. Here, we employ a computationally efficient model selection technique to map the relationship between the best-fit models – Beta-Poisson (Telegraph), NB, and Poisson – and the kinetic parameters that govern gene expression stochasticity. Our findings reveal that the NB distribution closely approximates simulated data (incorporating both biological and technical noise) within an intermediate range of the sum of the gene activation and inactivation rates normalized by the mRNA degradation rate. This range expands with decreasing mean expression, increasing technical noise, and larger sample sizes. The results imply that: (i) good NB fits occur in diverse parameter regimes without exclusively indicating transcriptional bursting; (ii) for small sample sizes, biological noise predominantly shapes the NB profile even when technical noise is present; (iii) under steady-state conditions, gene-specific parameters (burst size and frequency) estimated in regions where the NB model fits well, typically show large relative errors, even after corrections for technical noise, and (iv) gene ranking by burst frequency remains reliably accurate, suggesting that burst parameters are most informative in a relative sense. Finally, applying technical-noise–corrected model fitting to scRNA-seq data confirms that a substantial fraction of mammalian genes fall within these NB-fitting regimes, despite lacking transcriptional bursting.

## 1 Introduction

Single-cell RNA sequencing (scRNA-seq) facilitates the quantitative characterization of cellular heterogeneity by providing genome-wide transcriptomic profiles at single-cell resolution across hundreds or thousands of individual cells [1–4]. The use of unique molecular identifiers (UMIs) has substantially improved the reliability of sequencing data by enabling the distinction between original RNA molecules and PCR duplicates. By tagging each molecule with a unique barcode prior to amplification, UMIs mitigate amplification biases and sequencing artifacts, resulting in molecular counts that more accurately reflect true biological abundance [5]. The transcript count distribution of most genes is unimodal and well approximated by a negative binomial (NB) distribution, a two-parameter model often interpreted as a Gamma–Poisson mixture, where one parameter reflects the number of successes and the other the success probability [6–11]. The significance of the NB distribution is highlighted by its extensive application in various computational tools for scRNA-seq analysis [10, 12–19]. The fitting of the NB distribution to data yields two parameters for each gene, providing a convenient numerical signature of its observed stochasticity. However, the origin of the universality of the NB distribution in single-cell sequencing data remains unclear. In fact, this distribution is also commonly found using single-molecule fluorescence in situ hybridization (smFISH), suggesting that its ubiquity cannot be simply dismissed as stemming from the significant technical noise of sequencing technologies [20].

Over the past two decades, using fluorescence-based single-cell transcriptomics, it has been shown that biological noise for each gene is well described by the two-state telegraph model [21–27] or by a three or more-state extension of this model [28–30]. Here, we focus on the telegraph model because it has been shown to very well approximate the steady-state distribution of mRNA counts predicted by models with a higher number of states [31–33]. The telegraph model describes a gene that switches between an active state (*G*) and an inactive state (*G*^∗^). Transcription occurs only from the active state *G*, and subsequently mRNA decay occurs with first-order kinetics. For an illustration, see Fig. 1a. The solution of the chemical master equation that describes the telegraph model in steady-state conditions leads to a Beta-Poisson distribution of mRNA counts [34–36]. Typically, scRNA-seq measurements detect a significantly smaller fraction of transcripts than smFISH and hence extending the telegraph model to also account for technical noise is important. It has been shown that even after this modification, the steady-state distribution of transcript counts in a single cell follows a Beta-Poisson distribution [37, 38] — technical noise simply leads to a rescaling of the effective transcription rate by the probability of transcript capture in the cell, but does not change the type of distribution. Depending on the parameter values, the Beta-Poisson distribution can be unimodal with a peak at zero, unimodal with a peak at a non-zero value, or bimodal with peaks at zero and non-zero values [39]. All of these distribution shapes have been measured using smFISH [20, 40, 41]. Hence, clearly, the classical model of stochastic gene expression, even when extended to account for technical noise, does not generally predict an NB distribution. There are two known conditions under which the Beta-Poisson distribution reduces to an NB distribution.

**Figure 1:**
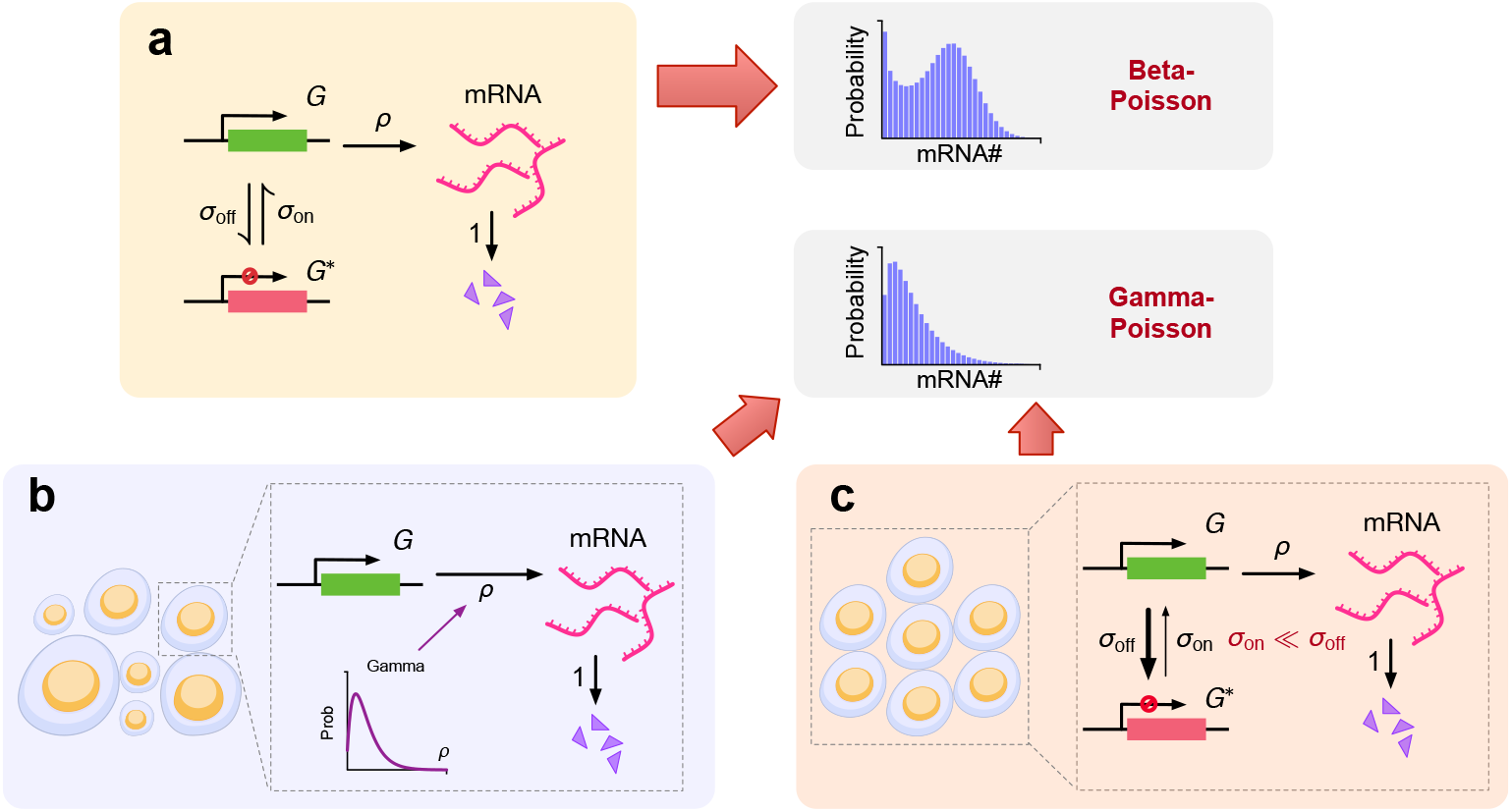
(a) Schematic illustrating the telegraph model of gene expression. A gene switches between active (green) and inactive states (red) with rates *σ*_on_ and *σ*_off_. Synthesis of transcripts occurs from the active state with rate *ρ*. The transcripts are subsequently degraded with rate 1. The rates are all normalized by the degradation rate. The steady-state distribution of transcript numbers is a Beta-Poisson compound distribution; (b) Schematic showing the special case where the gene is always in the active state and the transcription rate *ρ* varies from cell to cell according to a Gamma distribution. In this case, the mRNA distribution predicted by the telegraph model reduces to a Gamma-Poisson compound distribution (an NB distribution); (c) Schematic showing the special case where the gene spends most of its time in the inactive state (*σ*_on_ ≪ *σ*_off_) which leads to transcription occurring in short-lived bursts that are well separated from each other. All cells are identical, i.e. the rate constants do not vary from cell to cell. In this case, the mRNA distribution predicted by the telegraph model also reduces to an NB distribution.

Case (i). Say that a gene is always in the active state, i.e. there is no gene state switching. The telegraph model predicts that, in each cell, the transcript counts are sampled from a Poisson distribution with a parameter given by the effective transcription rate divided by the degradation rate [42]. If the effective transcription rate, which is equal to the product of the transcription rate and the probability of transcript capture, is the same in each cell, then the distribution of transcript counts across all cells is necessarily Poisson. However, if the effective transcription rate varies from cell to cell according to a Gamma distribution, then it immediately follows that the observed distribution is a Gamma-Poisson compound distribution, which is an NB distribution. Hence, in this case, the NB character of the distribution stems from extrinsic (cell-to-cell variation in the transcription rate) or technical noise (cell-to-cell variation in the transcript capture probability), and not from the intrinsic dynamics of a gene. For an illustration, see Fig. 1b. This case is unlikely to be the main reason for the universality of the NB in scRNA-seq data because the visualization of transcription in living cells using live-cell imaging conclusively shows alternating periods of gene activity and inactivity [43]. This is due to many factors such as reversible binding of transcription factors, enhancer-promoter interactions, the clustering dynamics of Pol II and the opening and closing of chromatin [44–48].

Case (ii). Now consider the case where the gene spends most of its time in the inactive state and synthesizes mRNA in a burst, i.e. in the short time that the gene is active. This occurs when the gene inactivation rate is much larger than the gene activation rate. This phenomenon is often referred to as transcriptional bursting [36]; for an illustration, see Fig. 1c. If the effective transcription rate is the same in each cell, then the Beta-Poisson distribution of the telegraph model has been shown to reduce to an NB distribution [49]. This case has sometimes been used to explain how the NB distribution of mRNA counts in sequencing data can arise from a physical model of gene expression [11]. However, this is also unlikely to explain the universality of the NB distribution in scRNA-seq data because the constraint that the activation rate is much lower than the inactivation rate represents a very small part of the available parameter space. Also because the assumption that the effective transcription rate is the same in all cells implies negligible variation in the probability of mRNA capture from one cell to another — which is difficult to reconcile with the wide distribution of total UMI counts (from all genes) per cell in typical scRNA-seq datasets [17, 50, 51].

In summary, there is no strong theoretical reason behind the universality of the NB distribution in scRNA-seq data. The two existing results in the literature on stochastic gene expression can only explain the NB distribution by invoking strong assumptions that are not realistic. Furthermore, it remains unclear what biological or biophysical meaning is to be imparted to the two parameters of the NB distribution. Currently, this meaning is only clear when gene expression is bursty (as in Case (ii)), in which case after correcting for technical noise, the two parameters can be used to extract the burst frequency (the rate at which mRNA bursts are transcribed) and the burst size (the mean number of mRNA produced when the gene is active) [52].

In this paper, we overcome these limitations by deriving new parametric conditions under which the observed distribution of transcript counts is well approximated by an NB distribution. We find that these conditions include, but are not limited to, transcriptional bursting. We also show that while the absolute values of the burst frequency and the burst size cannot generally be reliably extracted from the two parameters of the NB distribution, nevertheless these still contain useful information about differential gene expression.

## 2 Results

### 2.1 The NB distribution can provide a good approximation to the mRNA distribution of the telegraph model even when transcriptional bursting is absent

Consider an idealized scenario where each cell is identical to each other, i.e. for a given gene, there is no cell-to-cell variation in the parameters of the telegraph model (Fig. 1a). This implies that all cells are of the same type. Furthermore, assume that all transcripts from each cell can be detected. In this case, the distribution of mRNA counts is given by the steady-state solution of the chemical master equation of the telegraph model:

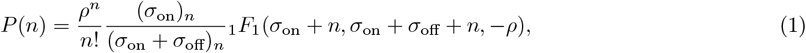

where *n* represents the mRNA count, 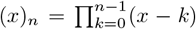 denotes the Pochhammer symbol, and 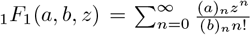 is the Kummer confluent hypergeometric function [34]. Note that *σ*_on_ is the gene activation rate, *σ*_off_ is the gene deactivation rate, *ρ* is the transcription rate; these rates are non-dimensional because they are normalized by the degradation rate (this convention will be used throughout the paper). Eq. (1) is equivalent to the compound Beta-Poisson distribution because the transcript numbers are distributed as follows:

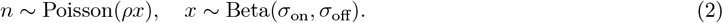

As mentioned previously, it is well known that if gene expression is bursty then Eq. (1) is well approximated by an NB distribution, or its continuous counterpart, the Gamma distribution [36, 49]; related derivations for the distribution of protein numbers can be found in Refs. [53, 54]. Specifically, transcriptional bursting occurs in the limit *σ*_off_ → ∞ taken such that *ρ/σ*_off_ remains constant [49, 55]. A simple way to see how this limit leads to an NB distribution makes use of the Beta-Poisson formulation of the telegraph model. From Eq. (2), it follows that

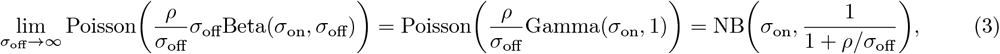

where in the second step we used the standard statistical result: 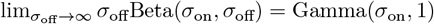.

#### It is presently unclear if the converse is true: Does an excellent NB fit to the mRNA count distribution of the telegraph model imply transcriptional bursting?

To answer this question, we proceed as follows. It is straightforward to show using Eq. (1) that the mean and variance of mRNA counts for the telegraph model are given by

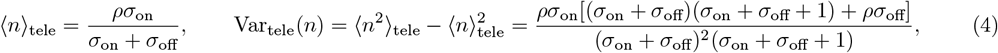

where ⟨•⟩ is the averaging operator. In contrast, the mean and variance of an NB distribution NB(*r, p*) are

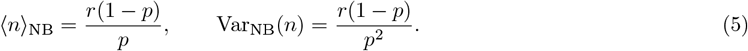

By equating ⟨*n* ⟩_tele_ = ⟨*n* ⟩_NB_ and Var_tele_(*n*) = Var_NB_(*n*) in Eqs. (4) and (5) and solving for *r* and *p*, we can construct an NB distribution that has the same first and second moments as the mRNA count distribution of the telegraph model:

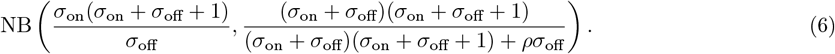

This distribution is hereafter referred to as the effective NB distribution denoted as NB(*r*_e_, *p*_e_). Of course, this distribution and the telegraph model potentially can differ in their third and higher moments, and hence there is no guarantee that the shapes of the two distributions will generally be similar. We also note that while maximum likelihood estimation or other methods can be used to obtain an effective negative binomial distribution, they often lack closed-form solutions for parameter estimates, which can hinder further quantitative analysis. In contrast, moment matching provides analytical expressions, facilitating deeper insight.

In Fig. 2, we compare the distributions given by Eq. (1) and Eq. (6) for four distinct parameter sets. In each case, the effective NB distribution is practically perfectly aligned with the corresponding telegraph distribution. However, note that only one of these four cases (Fig. 2a) corresponds to the classical case of *σ*_off_ ≫ *σ*_on_. The three counterexamples (Fig. 2b-2d) clearly show that excellent fitting of the telegraph model to an NB distribution does NOT necessarily indicate transcriptional bursting.

**Figure 2:**
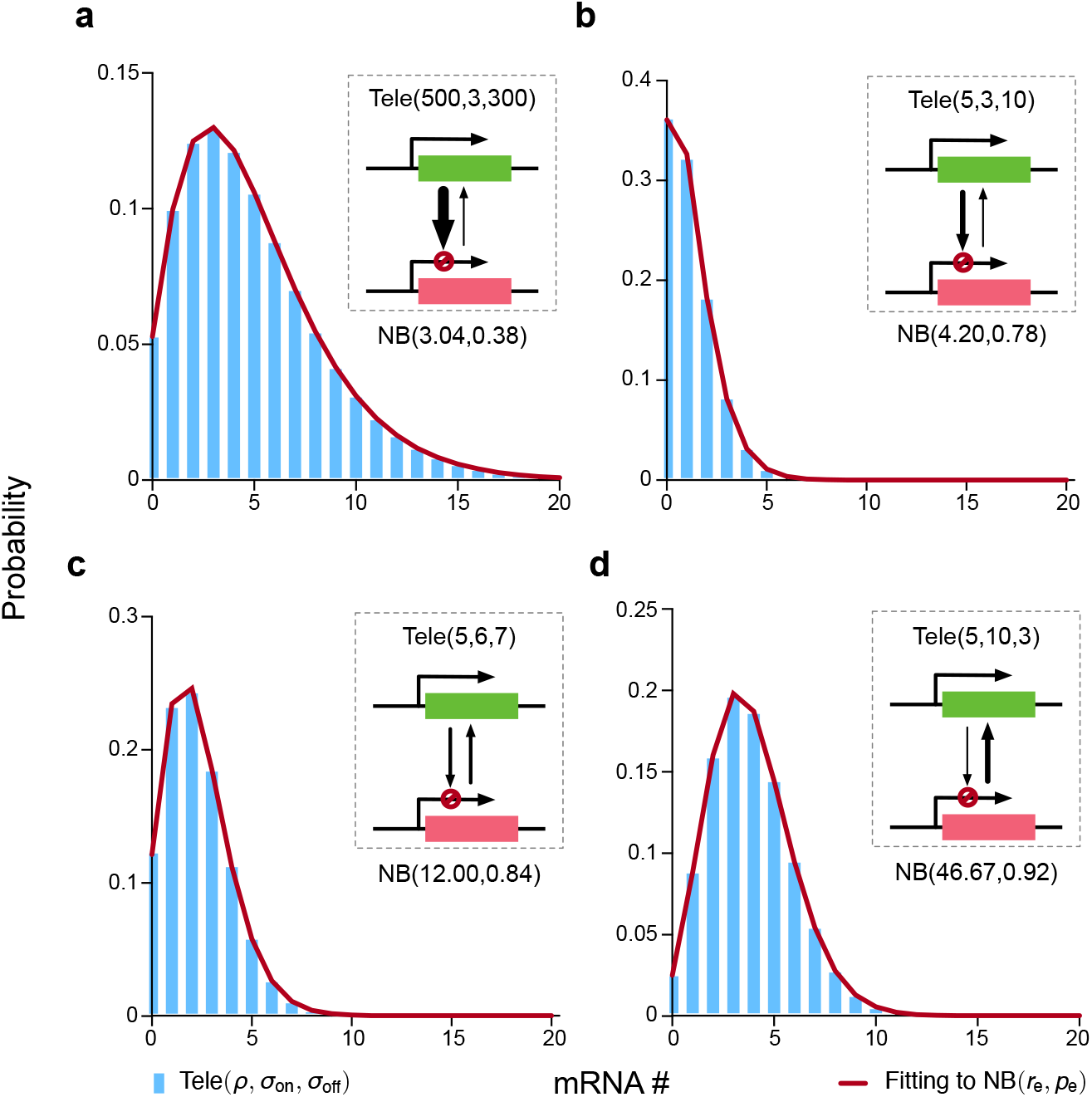
Comparison of the steady-state mRNA distribution of the telegraph model (Eq. (1) denoted as Tele(*ρ, σ*_on_, *σ*_off_)) with the effective NB distribution (Eq. (6) denoted as NB(*r*_e_, *p*_e_)) for the case of perfectly identical cells (parameters do not vary from cell-to-cell). The two parameters of the effective NB distribution are chosen so that its first and second moments of mRNA counts exactly agree with those of the telegraph model (the values of the two parameters, *r*_e_ and *p*_e_, are stated to 2 decimal places in the figure). In all four parameter cases, the effective NB distribution exceptionally well fits the corresponding telegraph distribution, and yet only in case (a) we have *σ*_on_ ≪ *σ*_off_ (the classical case of transcriptional bursting). This demonstrates that a good fit of an NB distribution to the telegraph model distribution does not imply the presence of transcriptional bursting.

### 2.2 An alternative set of conditions under which the NB distribution provides a good approximation to the telegraph model distribution

We start by reparameterizing the gene switching rates as

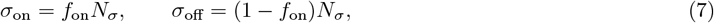

where *N*_*σ*_ = *σ*_on_ + *σ*_off_ represents the timescale of the transition rates relative to the rate of mRNA degradation and *f*_on_ is the fraction of time spent in the active state, which is given by

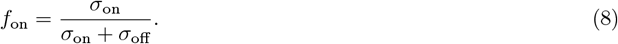

Furthermore, for convenience, we denote the telegraph model distribution, Eq. (1), by Tele(*ρ, σ*_on_, *σ*_off_). Given these definitions, we state the following theorem:

#### Theorem 1

In the limit *N*_*σ*_ → ∞, the telegraph model distribution Tele(*ρ, f*_on_*N*_*σ*_, (1 − *f*_on_)*N*_*σ*_) and the effective negative binomial distribution NB(*r*_e_, *p*_e_) (Eq. (6)) converge to the same distribution, both with a convergence rate of 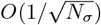.

The key idea behind proving this theorem is to show that the Beta and Gamma distributions converge to the same limiting distribution – specifically, a normal distribution – as *N*_*σ*_ → ∞. This then implies a convergence of the Beta-Poisson (telegraph model) distribution to a Gamma-Poisson (NB) distribution. This convergence is not immediately obvious because the Beta distribution is defined on (0, 1) while the Gamma distribution is defined on (0, ∞). Details of the proof can be found in Methods Section 4.1.

This proof makes more precise the statement in the caption of SI Fig. 4 of Ref. [56] which states that the mRNA count distribution of the telegraph model is bimodal (and therefore not similar to an NB distribution) when *σ*_on_ *<* 1 and *σ*_off_ *<* 1; it is unimodal and resembles an NB distribution when *σ*_on_ *>* 1 and *σ*_off_ *>* 1. In particular, Theorem 1 shows that *σ*_on_ *>* 1 and *σ*_off_ *>* 1 are not sufficient by itself to guarantee an excellent fit of the NB distribution to that of the telegraph model. We also note that the conditions identified by the theorem are different from those given by the classical result that requires *σ*_on_ ≪ *σ*_off_. In fact, while the classical results cannot explain the excellent fits shown in Fig. 2, these can be explained by Theorem 1, because in each of these cases *N*_*σ*_ is sufficiently large.

In the top left panel of Fig. 3, we use the Kullback–Leibler (KL) divergence, defined as 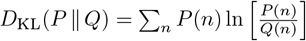 to quantify the proximity between the effective negative binomial distribution (*P* (*n*)) and the telegraph model distribution (*Q*(*n*)). For fixed values of *ρ* and *f*_on_, increasing *N*_*σ*_ results in a rapid decrease in the KL divergence, which agrees with Theorem 1. In the same figure, we also show the KL divergence between the Poisson distribution (with its parameter set to the mean of the telegraph model distribution) and the telegraph model distribution. It can be deduced that the fit of the Poisson distribution to the telegraph model distribution also becomes more accurate as *N*_*σ*_ increases, although it is always a poorer fit than the effective NB distribution. Of course, in the limit of large *N*_*σ*_, the KL divergence is very small for both effective NB and Poisson distributions and hence, practically speaking, in this limit both will provide an excellent approximation to the telegraph model distribution. These observations are further supported by comparing in Fig. 3 the mRNA count distributions predicted by the telegraph model with the effective NB distribution and the Poisson distribution for 5 different values of *N*_*σ*_ (these correspond to the points A-E in the plot of the KL divergence versus *N*_*σ*_ in the top left corner of Fig. 3).

**Table 1:** Parameter sets used in Fig. 4b and Fig. S1a,b.

|  | Parameters |  |
| --- | --- | --- |
| | $f_{\text{on}}$ | $N_{\sigma}$ |
| Set 1 | 0.1 | 1 |
| Set 2 | 0.4 | 10 |
| Set 3 | 0.2 | 10 |
| Set 4 | 0.7 | 100 |
| Set 5 | 0.5 | 1 |
| Set 6 | 0.9 | 0.1 |
| Set 7 | 0.3 | 1000 |
| Set 8 | 0.8 | 100 |
| Set 9 | 0.6 | 0.1 |
| Set 10 | 0.6 | 1000 |

**Figure 3:**
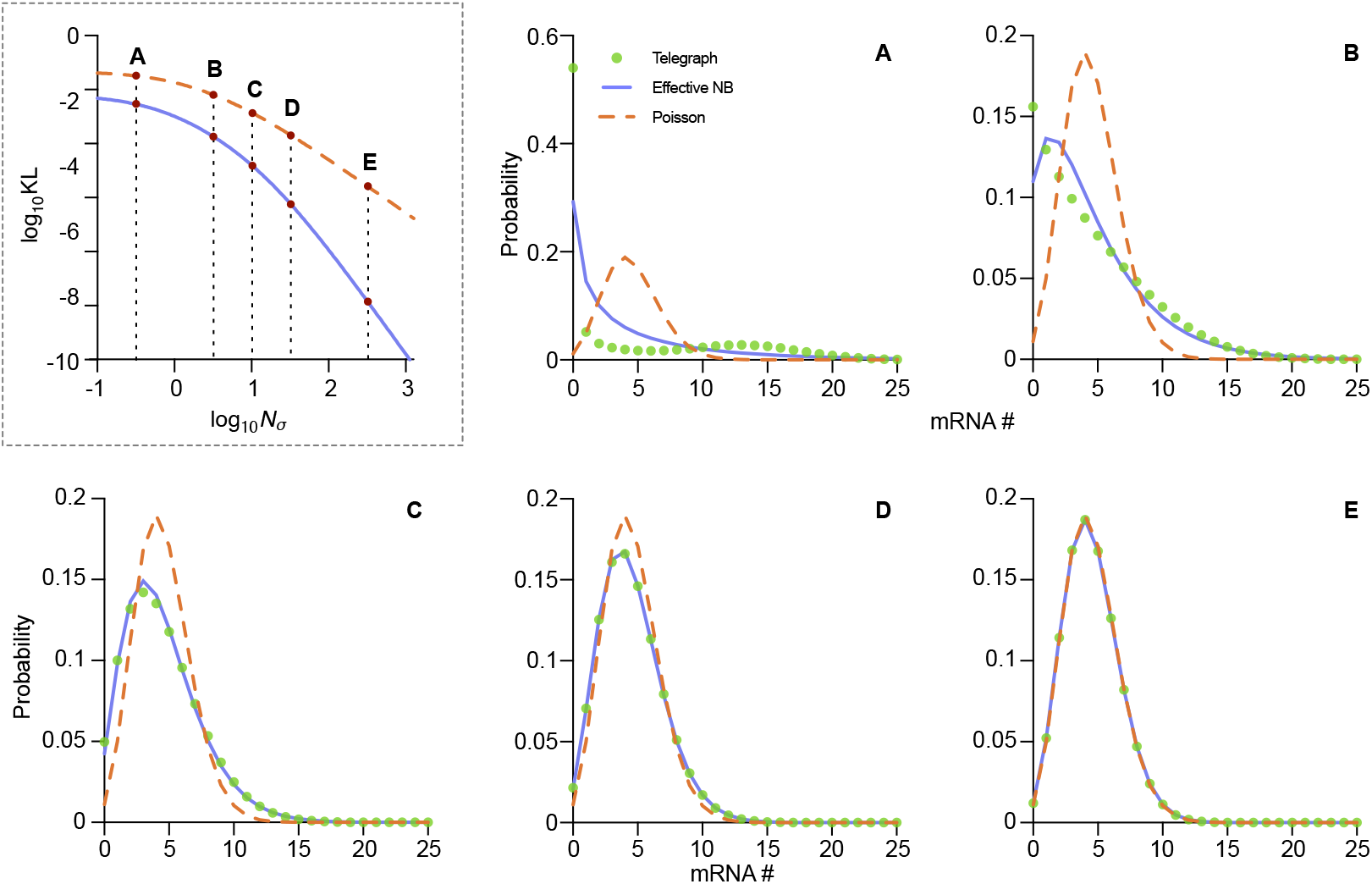
The mRNA distribution of the telegraph model converges to that of effective NB distribution as the sum of the gene-state switching rates relative to the mRNA degradation rate (*N*_*σ*_) increases. In the dashed box, we show that the effective NB distribution (blue solid line) exhibits a lower KL divergence to the telegraph model distribution compared to the Poisson approximation (orange dashed line) as *N*_*σ*_ grows. The distributions of the telegraph model (green dots), effective NB (blue solid lines), and Poisson (orange dashed lines) are shown for Points A-E, as indicated in the KL divergence plot. The other parameters are fixed at *ρ* = 15 and *f*_on_ = 0.3. Point A: *N*_*σ*_ = 0.3, Point B: *N*_*σ*_ = 3, Point C: *N*_*σ*_ = 10, Point D: *N*_*σ*_ = 30, Point E: *N*_*σ*_ = 300.

**Figure 4:**
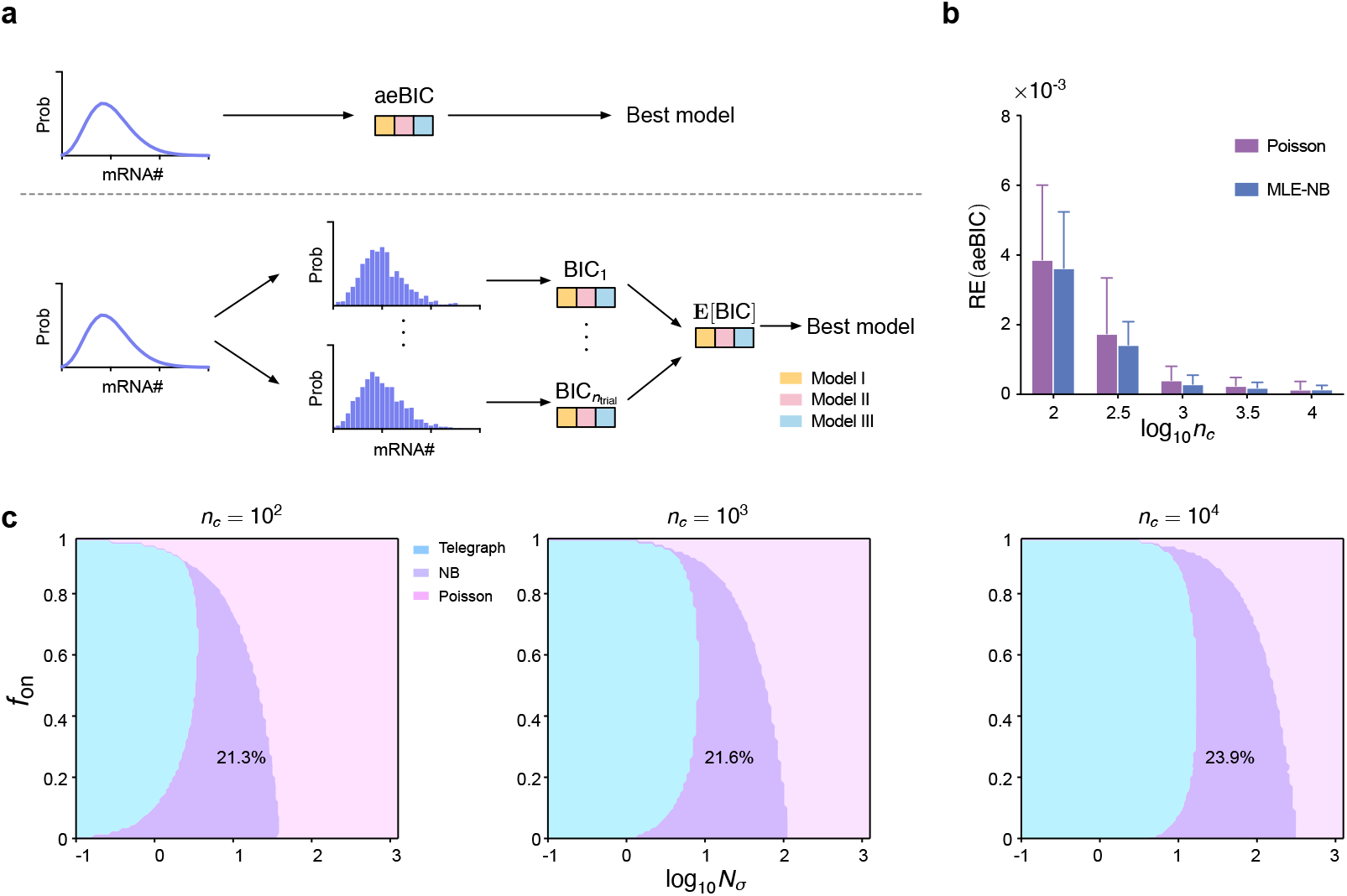
(a) Cartoon illustrating a computational approach to compare the aeBIC (top) with **E**[BIC] — the expected value of the BIC (bottom). The aeBIC utilizes a single score to select the best model distribution (telegraph, NB or Poisson) given that the ground-truth mRNA distribution is that of the telegraph model. The BIC method assigns a score to each different sample of simulated data from the telegraph model and then all these scores are averaged leading to **E**[BIC]. (b) The relative error (RE) of aeBIC compared to **E**[BIC] for two distributions (Poisson and NB) as a function of sample size *n*_*c*_ for 10 parameter sets (see Table 1 for the values of *N*_*σ*_ and *f*_on_; *ρ* is fixed to 15). Error bars show the standard error of the mean. (c) Phase diagram showing the regions of parameter space where the telegraph, NB and Poisson distributions are selected as optimal by the aeBIC, given that the ground-truth mRNA distribution is that of the telegraph model. Here *n*_*c*_ is the sample size, *N*_*σ*_ is the sum of gene-state switching rates normalised by the degradation rate of mRNA, and *f*_on_ is the fraction time spent in the active state. The fraction of the total parameter space occupied by the region where the NB distribution is optimally selected is shown on the plots. Note that the transcription rate is fixed to *ρ* = 15 which implies that the maximum mean number of transcripts in the phase plots is 15.

### 2.3 The NB distribution provides an optimal approximation to the telegraph model distribution for an intermediate range of the sum of the gene switching rates

The distribution comparison in Fig. 3 suggests that the effective NB distribution is an optimal fit to the telegraph model distribution in the intermediate range of *N*_*σ*_. This is because: (i) for small *N*_*σ*_, Theorem 1 does not hold. As well, in this case, it is already known that small *σ*_on_ and *σ*_off_ imply bimodal distributions of the telegraph model, which clearly cannot be well fitted by the (unimodal) effective NB distribution [56, 57]; (ii) for large *N*_*σ*_, the Poisson distribution practically provides an equally good fit as the effective NB distribution, but is preferable by the principle of Occam’s razor because it has one less parameter than the effective NB approximation.

To make this observation more rigorous, we will use a model selection approach. An often used method to select which of two models provides a best fit to the data involves (i) maximizing the likelihood function of each model on the sample data; (ii) evaluating the Bayesian Information Criterion (BIC) for each model which is a function of both the maximum value of the likelihood and the number of parameters; (iii) the optimal model is selected to be the one with the smallest BIC score.

In particular, the BIC is defined as

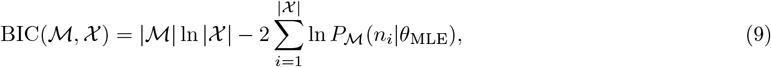

where 𝒳 represents the set of count data {*n*_*i*_} for *i* = 1, · · ·, *n*_*c*_, with |𝒳 | = *n*_*c*_ being the size of the dataset (number of cells). The proposed model is denoted by ℳ, its number of parameters by |ℳ| and the distribution that defines the model (its likelihood function) by *P*_ℳ_ (*n*_*i*_ | *θ*) where *θ* represents the model parameters. The parameter *θ*_MLE_ is the maximum likelihood estimate (MLE) obtained by maximizing the log-likelihood function

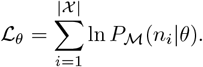

In our case, given a parameter set { *σ*_on_, *σ*_off_, *ρ* }, the count data is simulated by randomly sampling the telegraph model distribution *n*_*c*_ times. The difficulty with this approach is that for each parameter set, there is an infinite number of datasets that can be simulated, and in principle, the algorithm can select a different best model for each dataset. Rather than relying on individual samples, our goal is to associate a unique best model with each set of underlying parameters. The most straightforward way to achieve this is by averaging the BIC over an infinite number of samples, i.e., **E**_𝒳∼ 𝒢_ [BIC(ℳ, 𝒳)]. However, this approach is computationally intensive, as model parameters must be re-estimated for each sample.

To address this challenge, we introduce a new measure—the approximate expected Bayesian information criterion (aeBIC)—as an efficient estimator of the expected BIC across repeated samples. The aeBIC is defined as

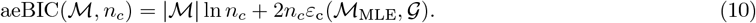

Here, *ε*_c_ represents the cross-entropy between two distributions (representing the model and the ground truth), defined as

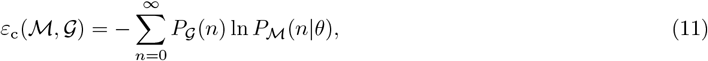

where *P*_𝒢_(*n*) and *P*_ℳ_(*n* | *θ*) are the probabilities of observing *n* molecules in distributions 𝒢 (the ground-truth distribution which for us is the telegraph model distribution) and ℳ (the proposed model which can be telegraph, NB or Poisson with parameters *θ*), respectively. Note that the subscript MLE denotes that the kinetic parameters of the proposed distribution are estimated via maximum likelihood. Specifically, ℳ_MLE_ represents the model whose parameters *θ* minimize Eq. (11). If ℳ= 𝒢, the cross entropy *ε*_c_(ℳ, 𝒢) reduces to the information entropy *ε*(𝒢). Additionally, the cross entropy can be decomposed as [58]

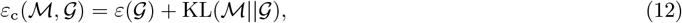

where KL(ℳ ||𝒢) is the KL divergence between ℳ and 𝒢. This decomposition implies that the closer the proposed distribution ℳ is to the ground-truth distribution 𝒢, the smaller the KL divergence becomes, and consequently, the cross entropy decreases as well.

In SI Sec 4.2 we show that the expectation of the BIC over an infinite number of independent samples (each of size *n*_*c*_) is related to the aeBIC by:

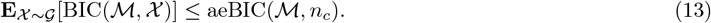

This shows that aeBIC serves as an upper bound for the expectation of the BIC. It can also be shown that the equality in Eq. (13) is achieved as *n*_*c*_ → ∞.

However, in practice, we find that the difference between **E**_𝒳∼𝒢_[BIC(ℳ, 𝒳)] and aeBIC(ℳ, *n*_*c*_) is negligible. To demonstrate this, for a fixed set of parameters { *ρ, σ*_on_, *σ*_off_ }, we randomly sampled *n*_*c*_ counts from the telegraph model distribution Tele(*ρ, σ*_on_, *σ*_off_) (using its Beta-Poisson formulation in Eq. (2)) and repeated this step *n*_trial_ = 10^3^ times. For each trial, we fitted the sampled counts to both Poisson and NB distributions using MLE and computed the BIC for both distributions. The BIC values were then averaged across all *n*_trial_ datasets to estimate the expectation of the BIC. Additionally, we computed the aeBIC using Eq. (10) for both distributions for each parameter set of the telegraph model distribution. The relative error of aeBIC compared to the expectation of BIC was calculated for each set of parameters. This comparison process is illustrated by a cartoon in Fig. 4a and the results for numerous parameter sets that cover the range *N*_*σ*_ ∈ [0.1, 1000], *f*_on_ ∈ [0.1, 0.9] and *ρ* ∈ [1, 15] are shown in Fig. 4b and Fig. S1a,b (these parameter sets correspond to a mean number of transcripts that ranges between 0 and 15). Note that, independent of the type of the fitted distribution, the relative error decreases with increasing sample size *n*_*c*_. Notably, even for small sample sizes, the magnitude of the relative error is very small, and thereby for all intents and purposes, we can equate the aeBIC with the average of the BIC computed over many independent samples. This is very convenient because it is far faster to compute the aeBIC than the BIC since, for the former, the maximum likelihood estimation only needs to be done once using Eq. (11).

Next, we use Eqs. (10)-(11) to compute the aeBIC and identify the best-fitted model (the one with the smallest aeBIC) among Poisson, NB and telegraph model distributions across the *N*_*σ*_ − *f*_on_ parameter space (while keeping the transcription rate *ρ* constant), given that the ground-truth data is generated by the telegraph model. Note that this model selection can be performed for different sample sizes because the aeBIC is a function of *n*_*c*_. Note also that the mean mRNA is proportional to *f*_on_ since the mean mRNA of the telegraph model is *ρf*_on_ (Eq. (4)) and *ρ* is fixed in our analysis. The results are shown in Fig. 4c — the model associated with the smallest aeBIC score corresponds to the model most frequently selected by the BIC calculated over many independent samples of the same size, thus providing a further test of the validity of the aeBIC approach (Table 2). In particular, aeBIC preferentially selects the telegraph model distribution for small *N*_*σ*_, while the Poisson distribution is preferred for large *N*_*σ*_. This agrees with the distribution comparison in Fig. 3 which suggested that the effective NB distribution is an optimal fit to the telegraph model distribution in an intermediate range of *N*_*σ*_. The NB distribution is more likely to be selected as *f*_on_ decreases, which is consistent with the fact that the telegraph model distribution converges to an NB distribution when the gene spends most of its time in the inactive state (*f*_on_ is small). Interestingly, the shape and area of the crescent-shaped region where the NB distribution is an optimal model remains largely unchanged as the sample size increases, but its position shifts horizontally. In particular, the fraction of space where the NB distribution is the optimal model increases monotonically from 21.3% to 30% as the sample size increases over four orders of magnitude, from 10^2^ to 10^6^ cells (results for 10^6^ cells are not shown).

**Table 2:** Comparison of the models selected by the aeBIC score and the BIC score for Fig. 4c with *n*_*c*_ = 100 cells. The minimum aeBIC score directly leads to a unique model that is shown in the first column. The BIC scores for the telegraph, NB and Poisson models are computed using MLE for each of 100 independent samples of size *n*_*c*_ generated by sampling from the Beta-Poisson formulation of the telegraph model Eq. (2) for the parameters shown in the 2nd and 3rd columns and *ρ* = 15; for each sample, the model with the smallest BIC score is selected (% of models selected are shown in the last 3 columns).

| Region | Parameters |  | Percentage of selected models by BIC |  |  |
| --- | --- | --- | --- | --- | --- |
| | $f_{\text{on}}$ | $N_\sigma$ | Telegraph | Negative Binomial | Poisson |
| Tele | 0.8 | 0.3 | 100% | 0% | 0% |
|  | 0.5 | 0.1 | 100% | 0% | 0% |
| NB | 0.4 | 10 | 7% | 88% | 5% |
|  | 0.2 | 7 | 7% | 93% | 0% |
| Pois | 0.9 | 100 | 0% | 1% | 99% |
|  | 0.1 | 400 | 0% | 1% | 99% |

Note that an alternative (and faster) way to obtain the phase space plots in Fig. 4c is to calculate the aeBIC with the value of the cross-entropy computed using moment matching between the proposed and the ground-truth distribution (instead of MLE). Specifically, the aeBIC using moment-matching is defined as

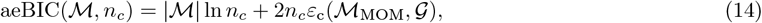

where *ε*_c_(ℳ_MOM_, 𝒢) is determined by evaluating Eq. (11) with parameters *θ* determined by moment-matching between the proposed distribution (*P*_ℳ_ (*n* | *θ*)) and the ground-truth distribution (*P*_𝒢_(*n*)). In Fig. S2 top left corner, we demonstrate that these two methods lead to very similar minimum values for the cross entropy and hence there is no appreciable difference on the model selection phase plots in Fig. 4c. On a MacBook Air with an Apple M2 chip and 16 𝒢B memory, generating a single phase space plot with this method (2.06 × 10^4^ pairs of *N*_*σ*_ − *f*_on_) takes about 68 seconds (CPU time) and uses only 3.6 MB of memory, demonstrating its computational efficiency.

### 2.4 Cell-to-cell variability in transcript capture probability significantly affects the region of parameter space where the NB distribution is optimally selected

Thus far, we have assumed that scRNA-seq data can be directly explained by the telegraph model. In practice, however, not all transcripts are captured. During the reverse transcription stage, each transcript is captured with probability *p*_1_. UMIs are attached to molecules before PCR, enabling PCR duplicates to be collapsed during downstream processing and thereby correcting amplification bias. Finally, during sequencing, the sequencing depth determines the detection probability (*p*_2_) of each UMI-labeled transcript. The observed UMI counts for a gene therefore represent the number of transcripts successfully captured, labeled, and detected. Collectively, the overall probability linking the true transcript count to the observed count is denoted as *p*_cap_ (including the effect of *p*_1_ and *p*_2_). Therefore, it is necessary to account for technical noise in the data. A simple way to do this is to assume that the number of zeros in the data is artificially high and to generate simulated scRNA-seq data using a zero-inflated telegraph model (a distribution that is a sum of a Dirac-delta function and the telegraph model distribution). While this approach or variants of it are widespread [14, 18, 19, 59] and attempts have been made to explain zero-inflation using mechanistic models [60], it is not ideal because (i) the downsampling of transcripts due to imperfect capture results in an increase of not only zeros but also 1’s, 2’s, etc; (ii) there is evidence that scRNA-seq data with UMIs barely suffer from zero-inflation [6, 61]. A more principled and increasingly used model to explain the effect of imperfect capture is the binomial capture model [10, 38, 62]. This model assumes that each transcript is captured and observed with a probability *p*_cap_ (Fig. 5a). For current standard droplet-based sequencing technologies, the capture probability typically ranges between 0.05 and 0.3, depending on the sequencing technique used [11, 63–65]. Extrinsic noise, such as variability in transcription rates, is a major contributor to cell-to-cell heterogeneity in gene expression [66, 67]. However, in most cases, variability in capture probability and transcription rate cannot be disentangled mathematically (see SI Section 4.3), unless additional experimental controls, such as spike-ins, are employed [8]. Therefore, in the following, we focus on variability in transcript capture probability.

**Figure 5:**
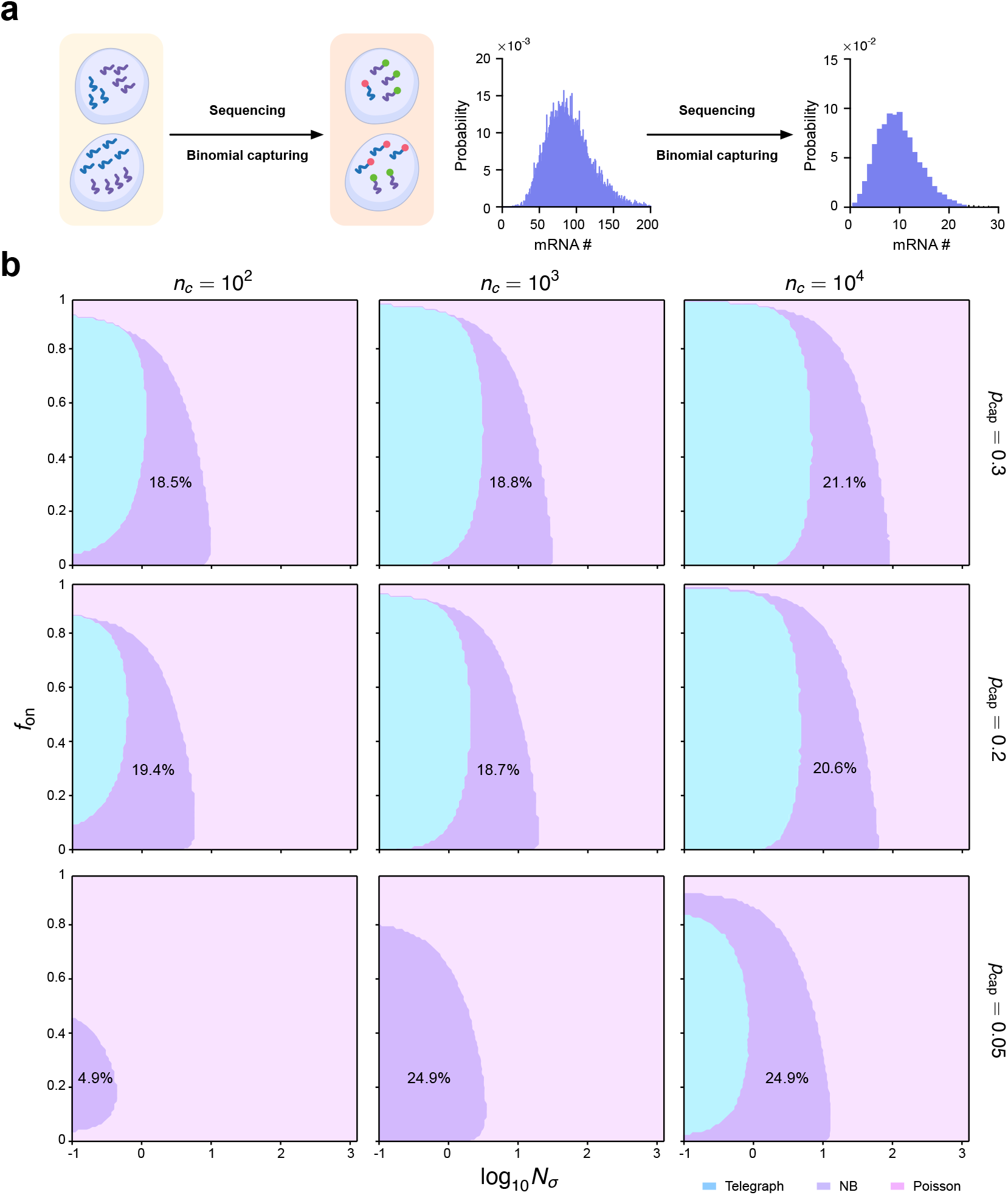
(a) Schematic illustrating the binomial capture model for scRNA-seq. Transcripts in each cell are captured with some probability *p*_cap_. This causes a downsampling of the distribution of mRNA counts. (b) Phase diagram showing the regions of parameter space where the telegraph, NB and Poisson distributions are selected as the optimal ones by the aeBIC, given that the ground-truth mRNA distribution is that of the telegraph model. The phase diagrams are shown for three different values of *p*_cap_ (values stated next to the plots). Here *n*_*c*_ is the sample size, *N*_*σ*_ is the sum of gene-state switching rates normalised by the degradation rate of mRNA, and *f*_on_ is the fraction time spent in the active state. The fraction of the total parameter space occupied by the region where the NB distribution is optimally selected is shown on the plots. Note that the transcription rate is fixed to *ρ* = 15 which implies that the maximum mean number of transcripts in the phase plots is 15*p*_cap_ = 4.5, 3 and 0.75 for the phase plots in rows 1, 2 and 3, respectively. These phase plots do not appreciably change if aeBIC is determined using moment-matching instead of MLE (Fig. S2).

For simplicity, first we consider the case where the capture probability does not vary from cell to cell and from gene to gene, i.e. the distribution of the capture probability is a Dirac-delta function, Dirac(*p*_cap_). We investigated the impact of *p*_cap_ on model selection outcomes (using aeBIC) and the results are shown in Fig. 5b. Note that in this case 𝒢 in Eq. (10) is the telegraph model distribution Eq.(1) with *ρ* rescaled to *ρp*_cap_ [37, 38]. Comparing Fig. 4c with Fig. 5b, we see that for a fixed sample size, the fraction of parameter space where the NB distribution is preferentially selected is almost constant for capture probabilities in the range *p*_cap_ = 0.2 − 1.0. The accuracy of scRNA-seq techniques is constantly improving, with commonly used platforms such as 10x having a capture probability that exceeds 0.2 [65]. Hence, we can conclude that if the capture probability is roughly the same from one cell to another, then the universality of the NB distribution in scRNA-seq datasets has little to do with the low capture efficiency.

Of course, for some scRNA-seq methods the capture probability may be very low and then in that case the technical noise will be so large that there will be a significant impact on model selection. For example, in the case of *p*_cap_ = 0.05 and sample size *n*_*c*_ = 10^2^ in Fig. 5b, the Poisson distribution emerges as the optimal model across most of the parameter space, with the NB distribution being selected only in a limited region. The telegraph model, however, is never identified as the optimal choice under these conditions. This is expected because the bimodal character of the telegraph model distribution at small *N*_*σ*_ becomes less obvious at lower capture probabilities — downsampling causes the two peaks of the distribution to become closer together or even to merge and hence the best-fit distributions are exclusively unimodal (Poisson or NB distributions).

The wide distribution of the total counts (from all genes) per cell in typical scRNA-seq datasets (see for e.g. Fig. 1A of [17]) strongly suggests that the capture probability varies considerably across cells (Fig. 6a). To address this, we simulate scRNA-seq data by sampling from the underlying mRNA distribution given by Eq. (2) with *ρ* rescaled to *ρp*_cap_, and *p*_cap_ that varies between cells according to three different distributions (Dirac-delta function and two different Beta distributions), all of which have mean ⟨ *p*_cap_ ⟩ = 0.3 but with different coefficients of variation (CV = 0 for the Dirac-delta function, and 0.11 and 0.21 for the two different Beta distributions). See Fig. 6b for a plot of the three distributions. Note that the Beta distribution was chosen because it samples numbers in the range (0, 1), a necessity given *p*_cap_ is a probability. The aeBIC was used to select between the standard telegraph model distribution (no assumption of variability of rates or capture probability between cells), NB and Poisson models with the ground-truth (measured) mRNA distribution 𝒢 given by that of the telegraph model with *p*_cap_ distributed according to a Beta distribution (SI section 4.4). Note that the aeBIC remains an accurate predictor of the expectation of the BIC computed from MLE and hence suitable for model selection even when the transcript numbers are heavily influenced by technical noise (Fig S1c). The phase plots generated by aeBIC-based model selection are shown in Fig. 6c. We find that for fixed sample size *n*_*c*_, an increase in the CV of the Beta distribution, leads to very small changes in the fraction of parameter space where the telegraph model distribution is selected, but the fraction of parameter space where the NB or Poisson distributions are selected changes significantly. In particular, an increase in the cell-to-cell variability in the capture probability tends to favor the selection of the NB distribution over the Poisson distribution. This is because variability in the capture probability increases the Fano factor (variance divided by the mean) of mRNA distributions (SI Section 4.5) — Fano factors larger than 1 can be captured by an NB distribution but the Poisson distribution has a fixed Fano factor of 1. For example, for a sample size of 10^3^ cells, as the CV of the Beta distribution of *p*_cap_ is increased from 0 to 0.21 (keeping the mean constant at 0.3), the fraction of space where the telegraph model distribution is optimal decreases slightly from 32.1% to 29.1%, where the NB distribution is optimal increases from 18.8% to 52.4%, and where the Poisson distribution is optimal decreases from 49.1% to 18.5%.

**Figure 6:**
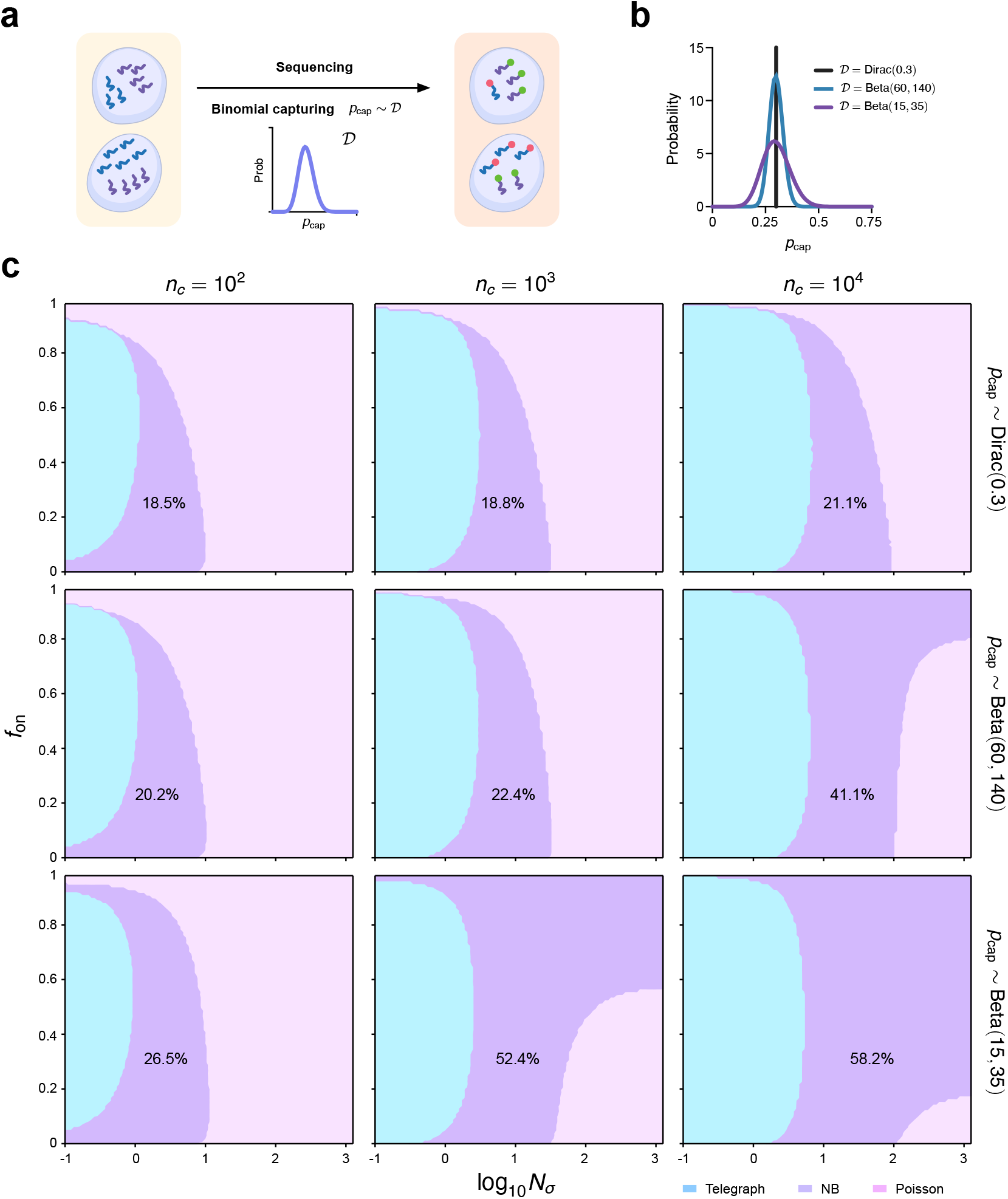
(a) Schematic illustrating the binomial capture model for scRNA-seq with a probability of mRNA capture, *p*_cap_, that varies between cells according to some distribution. (b) We consider three different distributions all with mean ⟨ *p*_cap_ ⟩ = 0.3 but with varying coefficient of variation (CV): (i) Dirac(0.3) with CV = 0; (ii) Beta(60, 140) with CV = 0.11; (iii) Beta(15, 35) with CV = 0.21. (c) Phase diagram showing the regions of parameter space where the telegraph, NB and Poisson distributions are selected as the optimal ones by the aeBIC, given that the ground-truth mRNA distribution is that of the telegraph model with effective transcription rate *ρp*_cap_ where *p*_cap_ is sampled from the 3 distributions mentioned above. Here *n*_*c*_ is the sample size, *N*_*σ*_ is the sum of gene-state switching rates normalised by the degradation rate of mRNA, and *f*_on_ is the fraction time spent in the active state. The fraction of the total parameter space occupied by the region where the NB distribution is optimally selected is shown on the plots. Note that the transcription rate is fixed to *ρ* = 15 which implies that the maximum mean number of transcripts in the phase plots is 15⟨*p*_cap_⟩ = 4.5.

### 2.5 In NB-optimal parameter space, burst parameter estimation accuracy is typically low, but ranking genes by burst frequency remains reliable

We also seek to understand if any biologically relevant interpretation can be imparted to the two parameters of the NB distribution fitted to the scRNA-seq data. Consider the ideal scenario in which we can perfectly correct for technical noise. In regions of parameter space where the NB distribution is optimally selected, can we obtain accurate estimates of the burst size and burst frequency from the two parameters of the best fitted NB distribution?

The aeBIC Eq. (10) used for model selection depends on the choice of the ground-truth model describing the distribution of the data (𝒢) and the model(s) (ℳ) to be fitted to this data. In our case, 𝒢 is the telegraph model with effective transcription rate *ρp*_cap_, and *p*_cap_ distributed according to a Beta distribution, Beta(*a, b*). Previously, we assumed that the models ℳ are given by the standard telegraph, NB and Poisson distributions. These models do not have any information on the distribution of *p*_cap_ and hence this is the case where model selection using aeBIC is done without correcting for technical noise. In contrast, a perfect correction for technical noise is possible if instead the models ℳare corrected for technical noise, i.e. the mRNA distributions of the telegraph model, NB and Poisson models are integrated over the Beta(*a, b*) distribution (this of course assumes perfect knowledge of the parameters *a* and *b* characterizing the technical noise). For a derivation of these distributions, see Section 4.4. The differences between the conventional models and those with *p*_cap_ sampled from a distribution are illustrated in Fig. 7a.

**Figure 7:**
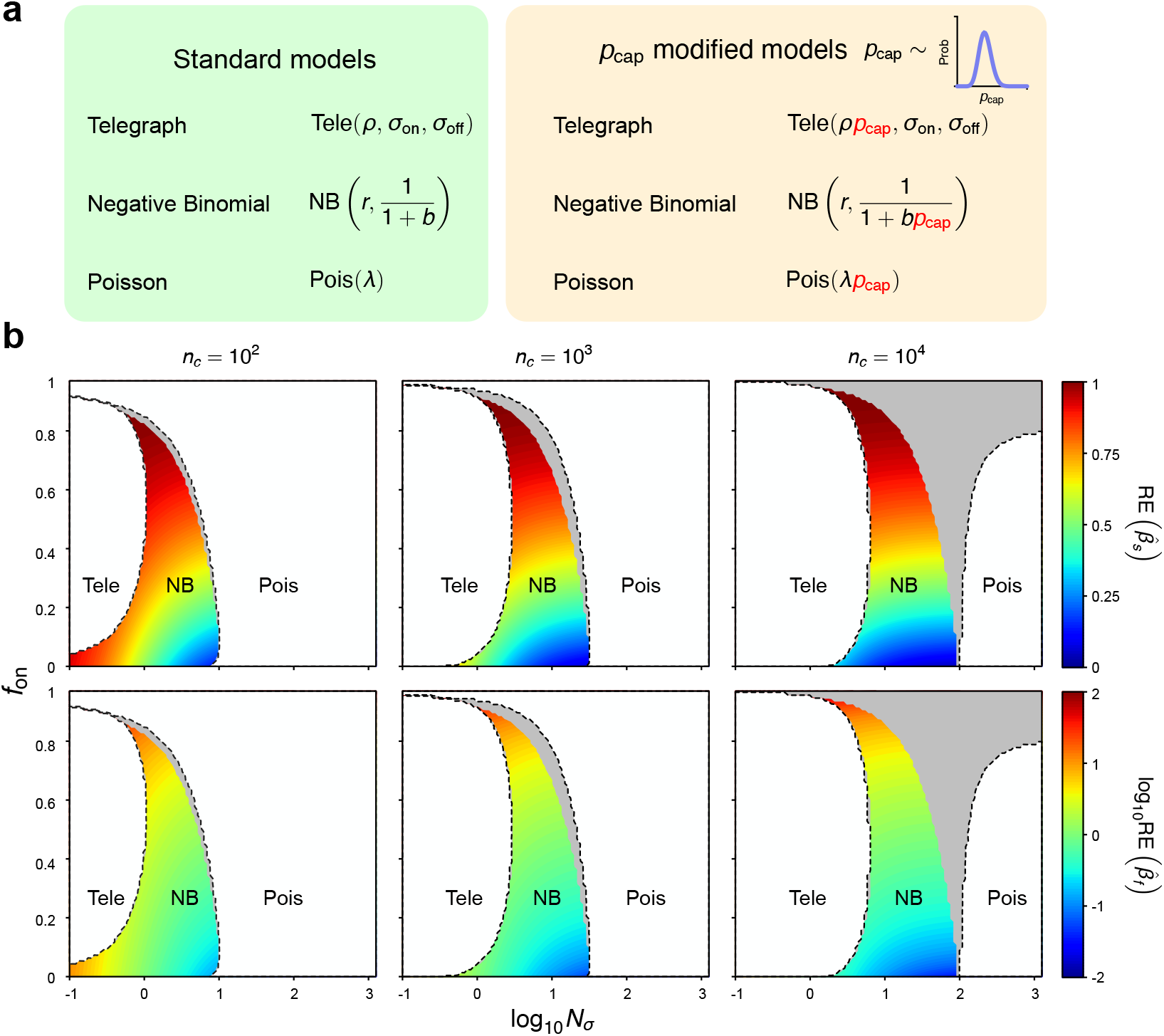
(a) Illustration of the differences between the standard and technical-noise-corrected models; (b) Phase diagrams produced by aeBIC model selection based on standard or corrected models. For both, the ground-truth model for observed data is the telegraph model with *p*_cap_ distributed according to the Beta(60, 140) distribution which has mean ⟨*p*_cap_ ⟩ = 0.3 and CV=0.11. The transcription rate *ρ* is fixed to 15; the maximum mean number of transcripts is 4.5. The labels “Tele”, “NB” and “Pois” denote the regions selected using the aeBIC procedure with corrected models. The dashed lines demarcate the same regions but using the aeBIC procedure with standard models. The “Pois” area is divided into a white part (where both aeBIC procedures select the Poisson distribution) and a grey part (where the aeBIC with standard models selects the NB distribution while the aeBIC with corrected models selects the Poisson distribution). The heatmap shows the magnitude of the relative errors in the estimated burst frequency 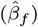 and burst size 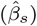 in the NB-optimal region (using the aeBIC with corrected models). The errors are computed using Eq. (17) — note that this approach assumes full knowledge of the distribution of probability capture, an ideal case. In the plots, *n*_*c*_ denotes sample size, *N*_*σ*_ is the sum of gene-state switching rates normalised by the degradation rate of mRNA, and *f*_on_ is the fraction time spent in the active state.

In Fig. 7b we contrast model selection using conventional and technical-noise-corrected models for the case where *p*_cap_ is distributed according to a Beta(60, 140) distribution which has a mean of ⟨ *p*_cap_ ⟩ = 0.3 and CV= 0.11. We find that the regions of parameter space where the conventional and corrected telegraph models are selected are very similar. However, the region where the corrected NB distribution is selected (rainbow-colored region) is smaller than the region where the conventional NB distribution is selected — in fact, there is a region (shown in grey) where the conventional NB distribution and the corrected Poisson distribution are selected, i.e. in this region the apparent NB character of the mRNA count distribution is purely due to technical noise. The discrepancies between the two approaches are negligible for small sample sizes (10^2^) but significant for large samples 10^4^.

Next, in the parameter space region where the corrected NB distribution is selected as the optimal model, we estimate the two parameters of the corrected NB distribution by the method of moments. We equate the first two moments of the latter distribution with those of the model simulating the data, i.e. the corrected telegraph model. This mimics the fitting of the corrected NB distribution to the data using the method of maximum likelihood in the limit of large sample sizes. In Section 4.5 we show that this procedure leads to NB(*r, p*) where *r* = *r*_e_ and *p* = *p*_e_ are exactly as given by the effective NB distribution in Eq. (6). Clearly, because *r*_e_ and *p*_e_ do not depend solely on the true burst frequency *σ*_on_ and burst size *ρ/σ*_off_ of the telegraph model [68], the latter two parameters cannot be directly estimated by moment matching.

This issue can be circumvented by interpreting the best-fitting corrected NB distribution NB(*r*_e_, *p*_e_) as equal to the steady-state NB distribution solution 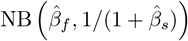 of the bursty gene expression circuit:

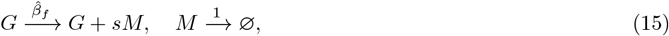

where *s* is a geometrically distributed random burst size with mean 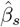 and the burst frequency is 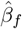 [69]. This model has experimental support [26] and its steady-state distribution is exactly the same as that of the telegraph model in the limit of transcriptional bursting, i.e. when the inactivation rate is much larger than the activation rate.

Hence the described inference procedure leads to estimated burst frequency and size that are given by

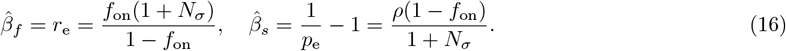

Since we know that the true burst frequency of the telegraph model is *σ*_on_ and the true burst size is *ρ/σ*_off_, it follows that the relative errors are given by

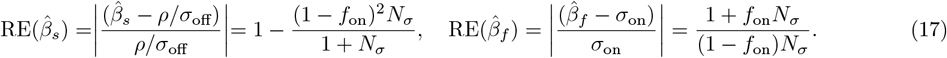

In Fig. 7b we show a heatmap of the relative errors in the NB-optimal region (using the aeBIC with technicalnoise-corrected models). The magnitude of these errors vary significantly and are smallest when *f*_on_ is small, i.e. when *σ*_off_ ≫ *σ*_on_ and *σ*_off_ ≫ 1. Hence, this implies that even though the NB distribution may be optimally selected, nevertheless this does not guarantee the accuracy of the estimated burst parameters. Indeed, the lack of accuracy is not very surprising given that the NB distribution of the reaction scheme in 15 is only a good approximation to the telegraph model distribution when *σ*_on_ ≪ *σ*_off_ [49] and that we have previously established that it is possible to have excellent NB fits even when this inequality does not hold (Fig. 2). Note that the burst size can generally be estimated more reliably than the burst frequency — this is also apparent from Eq. (17) which implies 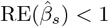 and 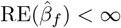.

Next, we sought to understand whether reliable information of some type can be extracted from the burst parameters, even though their absolute values are generally inaccurate. In particular, we seek to understand whether ranking of genes by their burst parameter values can be accurate since this is based on relative rather than absolute information. To answer this question, we used the following protocol:

1. Randomly sample two pairs of parameter sets 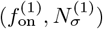 and 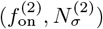, each associated with a different gene, from the parameter space region where the NB distribution (corrected for technical noise) is selected as the optimal model for *p*_cap_ distributed according to the Dirac (0, 3), Beta(60, 140) and Beta(15, 35) distributions. For a sample size of *n*_*c*_ = 10^4^, these regions are shown in purple in Fig. S3. Note that the transcription rate *ρ* is fixed to 15 in all cases.
2. Calculate the true burst frequency and size for each gene. For the first gene, we denote the true burst frequency and burst size as 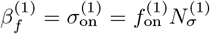 and 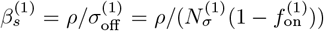, respectively; for the second gene, 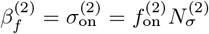 and 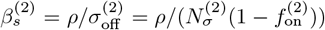, respectively. These relationships follow from Eq. (7).
3. For each gene, use the method of moments to fit an NB distribution integrated over the distribution of *p*_cap_ to the distribution of the data, i.e. telegraph model distribution integrated over the distribution of *p*_cap_. This leads to the estimated burst frequency-size pairs for each gene: 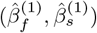 and 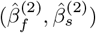 which are given by Eq. (16).
4. Compute the ground-truth ratios of burst sizes for the pair of genes as 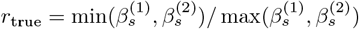. Note that by placing the smaller burst size in the numerator, we guarantee *r*_true_ *<* 1. The ground-truth ratio for the burst frequencies can be computed similarly.
5. Compute the estimated ratio of burst sizes for the pair of genes *r*_estimate_, preserving the original pair order used for the ground-truth ratio of burst sizes. That is, if 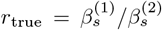 then 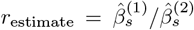 if 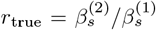 then 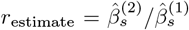.The estimated ratio for the burst frequencies can be computed similarly.

In Fig. 8a we show a plot of *r*_true_ versus *r*_estimate_. Each point in this scatter plot corresponds to a randomly selected pair of genes. Points are classified by their color: (i) blue if *r*_estimate_ *>* 1 indicates a reversal of pair order which implies that burst parameter inference fails to preserve the relative magnitude of the burst parameters; (ii) orange if *r*_estimate_ *<* 1 and *r*_estimate_ *> r*_true_ indicates that the order of the estimated burst parameters is the same as that of the ground truth parameters but that the estimation leads to an amplification of the difference between the burst parameters of the two genes; (iii) green if *r*_estimate_ *<* 1 and *r*_estimate_ *< r*_true_ indicates that the order of the estimated burst parameters is the same as that of the ground truth parameters but that the estimation leads to a reduction of the difference between the parameters of the two genes.

**Figure 8:**
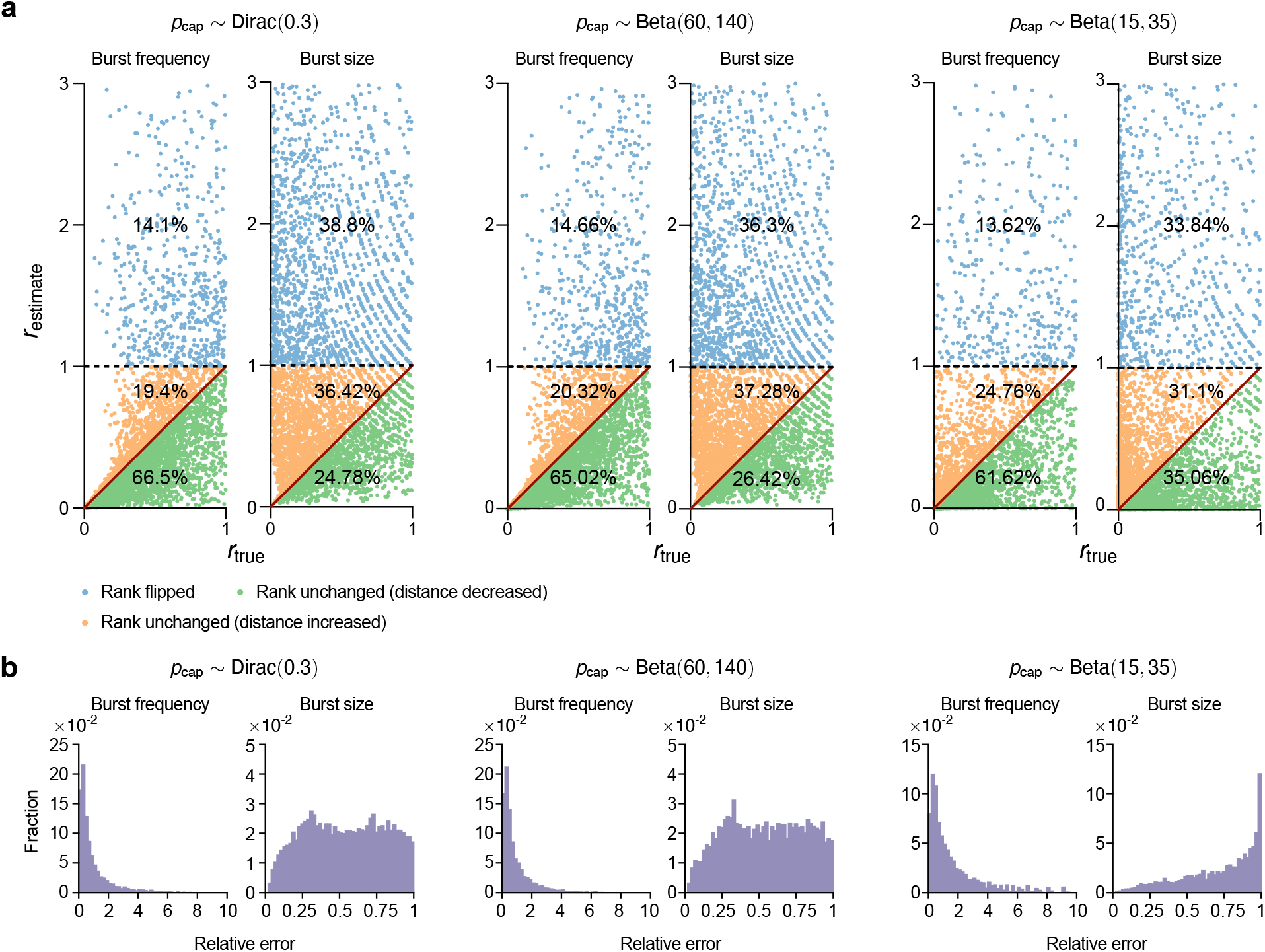
(a) Scatter plot of the estimated ratio of burst parameters *r*_estimate_ of a pair of genes and of the ground-truth ratio of the burst parameters *r*_true_ of the same pair of genes. All parameter sets are sampled from the region of parameter space where the technical-noise-corrected NB distribution is optimally selected using the aeBIC approach (purple regions in Fig. S3). Points marked as blue are those pairs of genes for which the estimation led to an incorrect ordering of genes by the size of the burst parameter. The order was correctly inferred for gene pairs corresponding to orange and green points. Perfect ratio estimation is shown by the solid red line; gene pairs corresponding to orange (green) points overestimate (underestimate) the distance between the burst parameters of the gene pairs. (b) Distributions of the relative errors in burst frequency and burst size for those pairs of genes for which the order was correctly inferred (orange and green points) in (a). Note that the inference and model selection approach here assumes full knowledge of the distribution of probability capture, an ideal case.

The results show that in many instances the ranking of a pair of genes by the size of their burst frequency is correctly estimated — these are the orange and green points in Fig. 8a which imply gene ranking accuracy by burst frequency in 86 − 87% of cases. However, the ranking of genes by burst size is significantly less accurate: it is correct in merely 58 − 69% of all cases, though it is better than expected by chance. In Fig. 8b we show the relative errors of the burst parameters for the pairs of genes that were correctly ranked (orange and green points in (a)). This verifies that generally relative errors and ranking accuracy are not related to each other since the ranking can be accurate and yet the relative errors can be large.

The main limitations of our approach are that we assume: (i) perfect knowledge of the distribution of capture probability; (ii) the information in the first two moments of the transcript count distribution is enough to accurately infer the burst parameters; (iii) we exclusively perform parameter inference using the technical-noise corrected NB model. To overcome these limitations, in Section 4.6, we describe an approach which simulates the workflow of an actual scRNA-seq experiment: (i) synthetic count data is generated for thousands of genes in *n*_*c*_ cells, which includes noise from both biological and technical sources; technical noise is modeled by a Beta distributed transcript capture probability; (ii) a maximum likelihood-based approach is used to infer the burst frequency and size, and to select between the technical-noise corrected telegraph, NB and Poisson models derived in Section 4.4. Note that this approach relies on using the measured total counts from all genes in a cell as a proxy for the (unknown) capture probability for that cell; the approach also does not simply use the first two moments of the observed distribution of counts, but the full distribution information and hence overcomes some of the known limitations of moment-matching approaches [70]; (iii) for those genes for which the technical-noise corrected NB model is selected as optimal by the BIC, the burst frequency and size are estimated separately from the technical-noise corrected telegraph and NB models. These are compared to the ground-truth values to calculate the relative errors. In addition, the percentage of pairs of genes that are correctly ranked according to the magnitude of their burst frequency and size, are calculated for each model. The whole methodology is illustrated in Fig. S4a. We considered three variations of this procedure, according to the generation of synthetic data in step (i):(a) *n*_*c*_ = 10^3^ cells, *N*_*σ*_ = 10 and the distribution of *p*_cap_ is fixed to Beta(15, 35); (b) *n*_*c*_ = 10^4^ cells, *N*_*σ*_ = 10 and the distribution of *p*_cap_ is fixed to Beta(60, 140); (c) *n*_*c*_ = 10^4^ cells, *N*_*σ*_ = 20 and the distribution of *p*_cap_ is fixed to Beta(60, 140).

The results for (a) are shown in Fig. S4 and for (b) and (c) are shown in Fig. S5. We find that for each fixed value of *N*_*σ*_, the relative errors of the burst frequency and size increase with the fraction of time that the gene spends in the active state *f*_on_. This agrees with the results shown in Fig. 7. Interestingly, even though the genes analyzed are those for which the technical-noise corrected NB model is selected as the optimal model by BIC, the parameter estimates are more accurate using the technical-noise corrected telegraph model (compare blue and green bars in Fig. S4(b), Fig. S5(a) and Fig. S5(c)). In addition, a high percentage of pairs of genes are correctly ranked according to the magnitude of their burst frequency, both by the technical noise-corrected NB and telegraph models; however, ranking by the burst size was much less reliable (see the piecharts in Fig. S4(c), Fig. S5(b) and Fig. S5(d))). The rankings also agree with the results shown in Fig. 8. Hence, overall, the results obtained from the MLE-based inference and BIC verify the accuracy of those previously obtained using moment-matching and aeBIC.

### 2.6 Analysis of experimental data reveals a substantial fraction of genes best fitted by NB but not transcriptionally bursty

Finally, we sought to determine how many genes that are best fitted by the negative binomial (NB) distribution are truly transcriptionally bursty, and whether this fraction is significant. Prior review papers, such as [71], provide ranges of switching rates (*σ*_on_ and *σ*_off_) that partially address this question. To answer it more comprehensively, we analyzed scRNA-seq data from mouse fibroblasts [72], preprocessing the dataset to ensure reliability (see SI Section 4.7). After preprocessing, 670 cells and 21,684 genes were retained.

We fitted the three *p*_cap_-modified models in Fig. 7a to this dataset using MLE. The distribution of *p*_cap_ was estimated via kernel density estimation, and the computation of count distributions over *p*_cap_ followed the procedure described in Section 4.8, consistent with the approach outlined in Supplementary Information Section VI of Ref. [73]. After parameter inference, we applied BIC to select the best-fitting model for each gene and found that ∼ 80% of genes were best fitted by the NB model (Fig. 9a).

**Figure 9:**
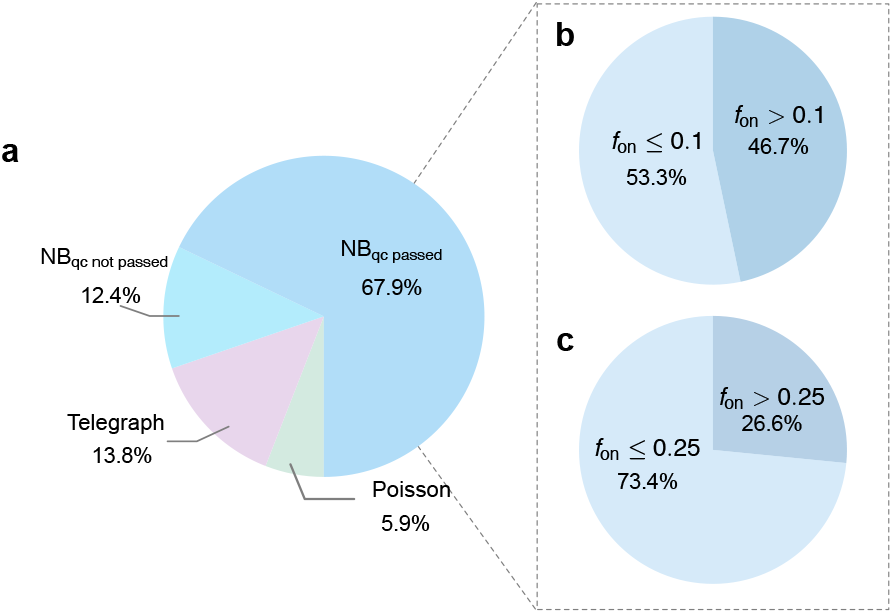
Analysis of mouse fibroblast scRNA-seq data reveals that many genes best fitted by the NB distribution are not transcriptionally bursty. (a) Model selection using BIC on 21,684 genes (670 cells) after preprocessing shows that ∼ 80% of genes are best fitted by the NB model. (b,c) For these NB-fitted genes, Bayesian inference of *f*_on_ was performed using the *p*_cap_-modified telegraph model, with reliability assessed by the confidence interval criterion (CI range/median *<* 0.4). The fraction of NB-fitted genes classified as transcriptionally bursty depends on the threshold chosen for *f*_on_: 0.1 in (b) and 0.25 in (c). In both cases, a substantial fraction of genes are best fitted by the NB model yet are not transcriptionally bursty.

For these NB-fitted genes, we refined the inference using Bayesian estimation with the `Turing.jl` package in Julia, fitting each gene’s count distribution with the *p*_cap_-modified telegraph model. We then quantified the confidence interval (CI) for *f*_on_. If the CI range (2.5%–97.5%) divided by the median was less than 0.4 (i.e., ± 20% around the median), we considered the estimate of *f*_on_ reliable. This yielded 14,722 genes, of which the inferred parameters have been summarized and deposited on GitHub (see Data availability statement).

We next computed the fraction of genes classified as transcriptionally bursty under different thresholds for *f*_on_ (0.1 in Fig. 9b and 0.25 in Fig. 9c). In both cases, a considerable fraction of genes were best fitted by the NB model yet not transcriptionally bursty. This result highlights the importance of our theoretical framework for correctly interpreting NB fits in scRNA-seq data.

## 3 Discussion

In this study, we have investigated why the NB distribution effectively approximates the transcript count distribution observed in scRNA-seq experiments and evaluated the biological implications of its two parameters. Presently, the universality of the NB distribution in UMI count data is puzzling given that a decade of analytical studies have unearthed only one stringent condition under which the distribution of the telegraph model of stochastic gene expression is well approximated by an NB distribution: genes must be predominantly inactive in a population of identical cells — conditions that are unlikely met in scRNA-seq experiments.

We first showed using theory and a new model selection criterion (aeBIC) that in the ideal case where the noise in the data is purely of biological origin, the NB distribution provides an accurate and optimal approximation of the transcript count distribution of a gene in a crescent-shaped region of its *f*_on_ vs *N*_*σ*_ parameter space where *f*_on_ is the fraction of time spent in the active state, while *N*_*σ*_ is the sum of the activation and deactivation rates (normalized by the degradation rate). This region occupied about 20% of the scanned parameter space, showing little dependence with the sample size (number of cells). In particular, the NB distribution is an excellent approximation in an intermediate range of *N*_*σ*_ and the size of this range increases with decreasing *f*_on_. The classical parameter regime where the NB distribution is thought to be a good approximation to the telegraph model, i.e. when *f*_on_ is small (genes mostly inactive) and transcriptional bursting is apparent [49], is a subset of the parameter regime that we have identified. In fact, this new region encompasses genes whose *f*_on_ ranges from small to large, thus showing that a good fit of the NB to the measured distribution of transcript counts is generally not indicative of transcriptional bursting.

Next, we investigated the more realistic case where the noise, i.e. the variability of mRNA numbers across cells, is due to both biological and technical noise. We simulated observed scRNA-seq data by downsampling the counts of the telegraph model of gene expression, assuming that the transcript capture probability for each cell is either constant or beta distributed. The selection of models through aeBIC allowed us to quantify how the proportion *F* of the parameter space where the NB distribution optimally fits the data varies with sample size and the shape of the capture probability distribution. We found that for a sample size of 100 cells where all noise is biological noise, *F* ≈ 21%; where technical noise is added assuming that each cell has a transcript capture probability of 0.3, *F* ≈ 19%; where technical noise is added assuming that the transcript capture probability is sampled from a Beta distribution with mean 0.3 and CV= 0.21, *F* ≈ 27%. In contrast, for a sample size of 10^4^ cells, the parameter space fractions for the aforementioned three cases were *F* ≈ 24%, *F* ≈ 21% and *F* ≈ 58%. Note that a mean capture probability of 0.3 is typical of 10x Genomics Chromium Single Cell 3’ version 3 reagent chemistry [65]. From these comparisons, we can deduce the following. If the capture probability distribution is: (i) narrow, the results are similar to those obtained under the assumption that all noise is biological; (ii) wide but the sample size is small (order 100 cells) then the results are also similar to those obtained under the assumption that all noise is biological; (iii) wide but for sample sizes of the order of thousands of cells, we find a significant enhancement of the region of parameter space where the NB distribution is the optimal model compared to the case where all noise is biological. Specifically, the proportion of the parameter space favoring the telegraph model remains largely constant, while the proportion favoring the NB distribution increases at the expense of the Poisson distribution. Hence, given that the capture probability distribution is wide in many cases (reflected by wide distributions of the total UMI counts per cell [17, 50, 51]), our results suggest that for small sample sizes, biological noise from those genes with an intermediate-sized *N*_*σ*_ parameter is the main determinant of the NB character of the count distribution, while for large sample sizes, technical noise leads to NB distributions in regions of parameter space where biological noise cannot.

A previous study by Tang et al. [37] generated a simulated data set consisting of 500 cells and 7000 genes by sampling from the steady-state distribution of the telegraph model (Eq. (1)). For each gene, the Akaike Information Criterion (AIC) scores were calculated for the telegraph, NB, and Poisson models, and the model with the lowest AIC was selected as the optimal fit. The set of genes for which the Poisson and NB models were favored tended to have a lower mean expression; however, the distributions of the means of the genes in the three groups (see Fig. 3a of [37]) overlapped significantly, indicating that the mean alone is not enough to explain the model that best fits the expression of each gene. In the phase diagrams in Fig. 4c we show the regions where the three models are selected as a function of *f*_on_ and *N*_*σ*_ — clearly *N*_*σ*_, not *f*_on_, is the primary determinant of the chosen model. Since the mean mRNA is proportional to *f*_on_ (the mean mRNA of the telegraph model is *ρf*_on_ and *ρ* is fixed in these phase plots), it follows that the selection of the optimal model is only weakly determined by the mean RNA. This explains why Tang et al. [37] did not observe a strong relationship between the selected model and the mean mRNA count. Another study [74] has also shown that the NB distribution is often complex enough to describe the simulated scRNA-seq data generated from the telegraph model but did not map the relationship between the choice of the best-fit model and the regions of parameter space, or account for technical noise, as we have done here. The complete mapping that we have reported was only possible due to the use of the novel model selection criterion, aeBIC, which is a computationally efficient and accurate estimator of the model most frequently selected by the standard maximum likelihood procedure followed by BIC.

We also investigated the reliability of the burst frequency and burst size estimated from the two parameters of the best-fitting NB distribution in the parameter space region where this model is selected as the optimal one. These are commonly estimated by interpreting the NB distribution as that arising from a simpler mechanistic model than the telegraph model, namely the bursty gene expression model (with reaction scheme Eq. (15)), which was first studied in [69] and is now commonly used as the basis for more sophisticated stochastic models of gene expression [75–80] including those used to fit scRNA-seq data [6, 52, 81–84]. This model can be derived from the telegraph model under the assumption that the gene inactivation rate is much larger than the gene activation rates [49], i.e. the transcriptional bursting regime. However, our analysis shows that when the parameter *N*_*σ*_ is large, the steady-state distribution of the telegraph model is well approximated by the negative binomial (NB) distribution, a scenario that applies beyond the bursting regime. This means that estimating the burst frequency and size of a gene by fitting the NB distribution solution of the bursty model Eq. (15) to scRNA-seq measurements of its expression (after correcting for technical noise) can lead to potentially large errors, even if the data is excellently fit by a NB distribution. Our analysis supports this notion — we find that in the region of parameter space where the distribution of the telegraph model is best fit by an NB distribution, the errors in the estimated burst frequency and size are generally large, easily exceeding 50% for the burst frequency and at most 100% for the burst size. However, interestingly, we observe that the relative magnitudes of the burst parameters still hold some validity (as also suggested by a recent study of the cell-cycle dependence of bursty gene expression [6]). In particular, we find that the ranking of two genes by their burst frequency estimate is correct in about 86 − 87% of the cases; for the burst size, the ranking is correct in 58 − 69% of the cases, showing less reliability. We also found that in the parameter space region where the NB model is preferentially selected, the limitations discussed above remain largely the same if instead one estimates the burst size and burst frequency from the three parameters obtained by fitting the distribution of the telegraph model (after correcting for technical noise) to the data using the method of maximum likelihood [35, 37, 85–88].

Our study also has some limitations. We have accounted for cell-to-cell variability in transcript counts due to intrinsic noise and differences in the effective transcription rate from one cell to another due to a varying transcript capture probability. Any variation in the effective transcription rate due to variability in the transcription rate (extrinsic noise on the transcription rate) between cells is indistinguishable from variability in the transcript capture probability and hence is automatically accounted for in our present method (see SI section 4.3). However, we have assumed that there is no cell-to-cell variation in the activation and inactivation rates, which is, of course, a simplification of biological reality. Within our phase diagram methodology, these effects can be properly incorporated using a ground-truth distribution 𝒢 in Eq. (10) that is derived via an integration of the distribution of the telegraph model over the distributions of the activation and inactivation rates — these are typically unknown but could be approximated by gamma or lognormal distributions as in [52]. The issues we have identified regarding the inaccuracy of the absolute values of the burst parameters are possibly because we have exclusively used the steady-state distributions of stochastic gene expression models. These are most commonly used for two reasons: (i) scRNA-seq data often come from a single snapshot measurement; (ii) it is much easier to solve the chemical master equation of gene expression models in steady-state conditions than in time [89]. However, the median mRNA half-life in mammalian cells is a sizeable fraction of the cell-cycle duration (for e.g. ≈ 7 hours [90] compared to ≈ 13 hours [91] in mouse embryonic stem cells), and hence it is unlikely that a steady state assumption generally holds in this case. To overcome this issue, one can first use methods like DeepCycle [92] or VeloCycle [93] to assign a cell age *θ* (which varies between 0 at the beginning of the cell-cycle and 1 at cell division) to each cell and then fit an age-dependent model of gene expression to the age-resolved scRNA-seq data [6]. Alternative ways to possibly circumvent the issues we have reported using steady-state models is to instead fit models that account for different cell states due to differentiation [94] or models that use a variety of single-cell data (transcriptomic, proteomic and epigenomic) [95]. These approaches are computationally challenging and are actively under investigation.

Additionally, the eigenvalues of the Hessian matrix of the log-likelihood quantify the curvature of the likelihood surface and reflect the degree of parameter degeneracy. As the telegraph model converges to the NB and further to the Poisson model with increasing *N*_*σ*_, the number of effective parameters decreases from three to two and finally to one. This reduction leads to broader confidence intervals for inferred parameters, with eigenvalue ratios diverging to infinity. However, as shown in Fig. S6, these ratios increase only gradually with *N*_*σ*_ and do not provide clear thresholds for model selection. In contrast, the aeBIC method yields well-defined selection boundaries, demonstrating that Hessian eigenvalues are not reliable indicators for model selection.

In conclusion, our findings establish that the NB distribution is a robust approximation of scRNA-seq transcript count distributions across diverse noise conditions and parameter regimes. Notably, its broad applicability challenges the conventional theoretical link to bursty gene expression, since both biological and technical noise can induce NB-like behavior even outside of classical conditions. Our analysis cautions against direct burst parameter estimation from NB or telegraph model fits and calls for nuanced model selection and parameter estimation approaches in single-cell genomics.

## Data Availability Statement

The code for this paper and the inferred parameters for Fig. 9 are available at https://github.com/Vivian0911unique/aeBIC.

## Acknowledgements

Z.C. acknowledges support from a Shanghai Action Plan for Technological Innovation Grant (23S41900500) and a Natural Science and Engineering Research Council of Canada Discovery Grant (RGPIN-2024-06015). R.G. acknowledges support from the Leverhulme Trust (RPG-2024-082). The authors are grateful to Augustinas Sukys for his valuable feedback on previous drafts of this work.

## Author contributions

Conceptualisation, Z.C. and R.G.; Methodology, Z.C. and R.G.; Software, Y.W. and Z.S.; Analysis, Y.W. and Z.S.; Visualisation, Y.W. and Z.S.; Writing – Original Draft, Z.C. and R.G.; Writing – Review & Editing, Z.C. and R.G.; Supervision, Z.C. and R.G.

## 4 Methods

### 4.1 Proof of Theorem 1

It is well known that the probability distribution of the Beta-Poisson process

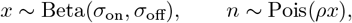

is identical to the steady-state distribution of the telegraph model given by Eq. (1) [35]. Another standard result is that the probability distribution of the Gamma-Poisson process

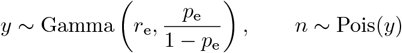

is identical to that of a negative binomial distribution NB(*r*_e_, *p*_e_). In what follows we shall fix these two parameters to be

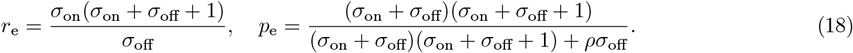

Note also that the definition of the Gamma distribution, Gamma(*α, β*), that we use is 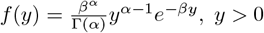, *y >* 0. Proposition 7 in Ref. [96] states that for two mixed Poisson distributions, denoted as MP(*g*_1_) and MP(*g*_2_), the convergence MP(*g*_1_) → MP(*g*_2_) holds if and only if *g*_1_ → *g*_2_ where → denotes convergence in distribution. Consequently, to prove Theorem 1, it suffices to demonstrate that the distribution of the random variable *x* converges to that of the random variable *y/ρ* as *N*_*σ*_ = *σ*_on_ + *σ*_off_ → ∞.

To proceed, we first determine the density function of the transformed random variable 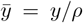. This is computed as

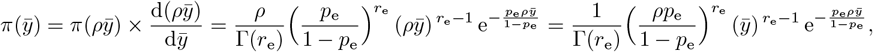

which precisely corresponds to the density function of 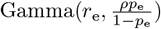.

To show 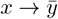, we need to establish the convergence

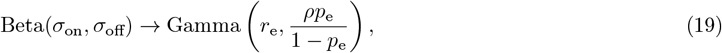

as *N*_*σ*_ = *σ*_on_ + *σ*_off_ → ∞. Rather than directly proving Eq. (19), we instead demonstrate that both distributions converge to a common limiting distribution, denoted as Dist, i.e.,

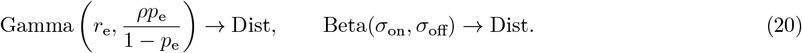

It turns out that Dist is a normal distribution.

Next, we show that the Gamma distribution in Eq. (20) converges to a normal distribution as *N*_*σ*_ → ∞. To achieve this, we introduce the moment generating function (MGF) for the random variable 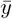,

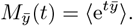

Consider the linear transformation

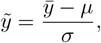

where

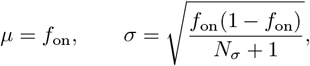

and 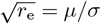. The MGF of the transformed random variable 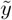 is

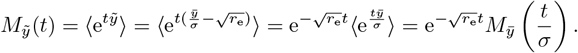

Using the MGF of the Gamma distribution, this simplifies to

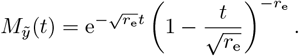

As *N*_*σ*_ → ∞, *r*_e_ → ∞, the expression further reduces to

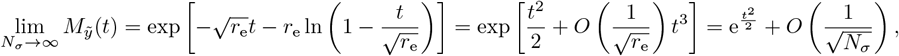

which is the MGF of the standard normal distribution 𝒩 (0, 1). Therefore, we conclude that

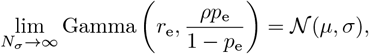

with convergence rate 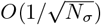.

Finally, we show that Beta(*σ*_on_, *σ*_off_) = Beta(*f*_on_*N*_*σ*_, (1 − *f*_on_)*N*_*σ*_) → 𝒩 (*µ, σ*) as *N*_*σ*_ → ∞. Note that we use the definition of the Beta distribution, Beta(*α, β*), given by

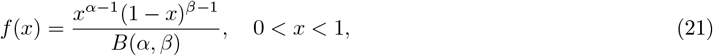

where *B*(*α, β*) = Γ(*α*)Γ(*β*)*/*Γ(*α* + *β*). The density function of this Beta distribution at the point *x* = *µ* + *σt* is given by

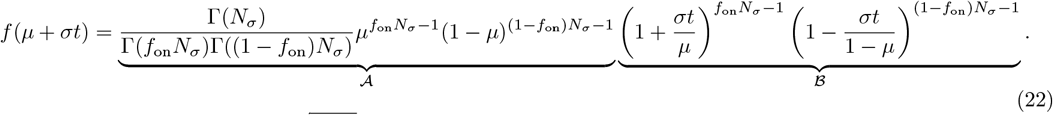

Using Stirling’s formula, 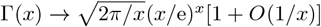 as *x* → ∞, the term 𝒜 in Eq. (22) reduces to

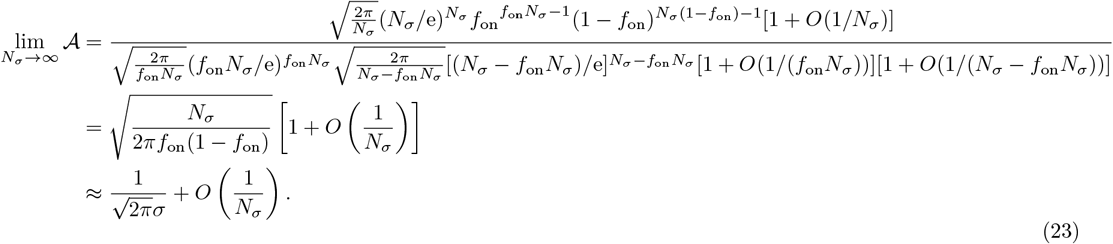

Next, we rewrite the term ℬ in Eq. (22) as

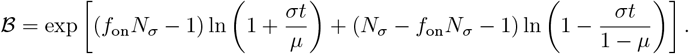

Noting that *σ/µ*→ 0 and *σ/*(1 − *µ*) → 0 as *N*_*σ*_ →∞, we expand ℬ around *t* = 0 using a Taylor series expansion, yielding

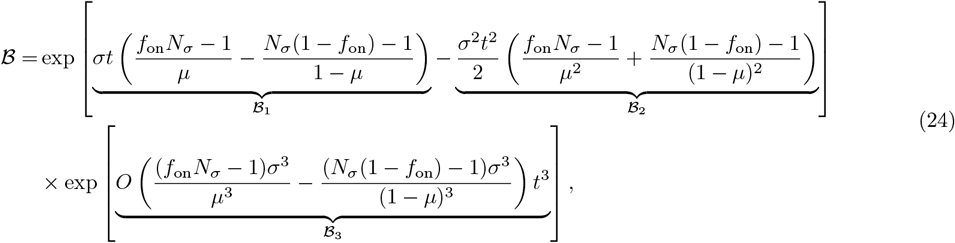

where

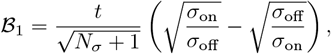

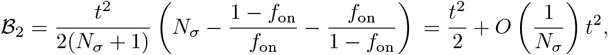

and

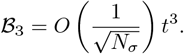

Using Eqs. (22)–(24), we obtain

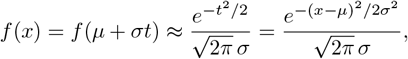

which corresponds to the probability density function of the normal distribution 𝒩 (*µ, σ*), with a convergence rate of 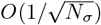. This concludes the proof of Theorem 1.

### 4.2 Mathematical relationship between aeBIC and BIC

Since the data in set 𝒳 (the set of count data {*n*_*i*_} for *i* = 1, · · ·, *n*_*c*_) are randomly sampled from the ground-truth distribution 𝒢, the expectation of the BIC (Eq. (9)) is given by

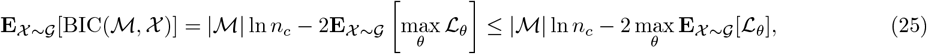

where the inequality follows from the fact that the expectation of the maximum is greater than or equal to the maximum of the expectation. A proof of this inequality is as follows.

For any fixed *θ*, it is true that

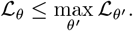

So taking expectations on both sides yields

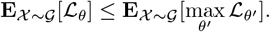

Because this is true for any *θ*, it follows that

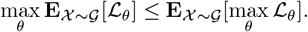

Furthermore, we note that

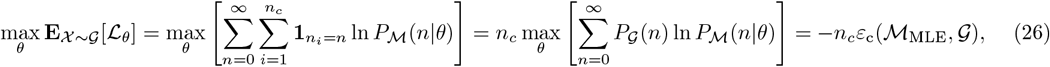

according to Eq. (11). Note that the first step in Eq. (26) is a reformulation of the log-likelihood function: the term 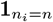 is an indicator function equal to 1 if and only if *n*_*i*_ = *n*, and hence 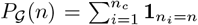. From Eqs. (10), (25), and (26), it follows that

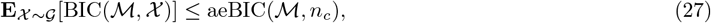

indicating that aeBIC serves as an upper bound for the expectation of the BIC. For a fixed distribution ℳ with fixed parameters, ℒ_*θ*_ → −*n*_*c*_*ε*_c_(ℳ, 𝒢) as *n*_*c*_ → ∞, which further suggests

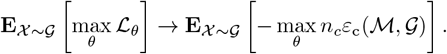

By noting that

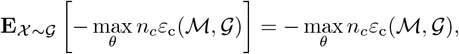

one can conclude that

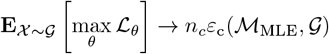

as *n*_*c*_ → ∞.

### 4.3 Breakdown of variability involved in transcription

Variability in scRNA-seq data arises from three major sources: intrinsic noise, extrinsic noise, and technical noise. The first two correspond to biological variability, whereas the latter is introduced by sequencing procedures.

**Intrinsic noise** mainly reflects stochastic promoter and chromatin activity, which is well captured by the gene-state switching dynamics of the telegraph model, as supported experimentally in [67]. The probability generating function (PGF) of the steady-state mRNA distribution under this model is

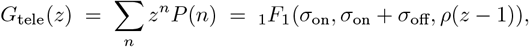

where *P* (*n*) is given in Eq. (1).

**Extrinsic noise** is primarily attributed to cell-to-cell variation in transcription rates [66, 67], manifested as heterogeneity in *ρ*.

**Technical noise** arises during sequencing, most notably from variability in transcript capture efficiency. Assuming each transcript is independently captured with probability *p*_cap_ [10, 62], the observed counts in each cell follow the PGF

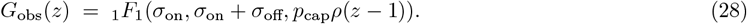

This result follows from the general property that binomial downsampling of a distribution with PGF *G*(*z*) produces a new PGF *G*(1 − *p*_cap_(1 − *z*)) (see [49], SI p. 9 for the special case *p*_cap_ = 1*/*2; for a more general discussion see [97]).

Importantly, Eq. (28) shows that the cell-specific transcription rate *ρ* and capture probability *p*_cap_ always appear as a product, *λ* = *ρp*_cap_, whose distribution can in principle be calculated from knowledge of the joint distribution *p*(*ρ, p*_cap_). This implies that, at the level of transcript count distributions, variability in transcription rates and variability in capture probability are mathematically indistinguishable. For this reason, in the following we focus our discussion on variability in transcript capture probability.

### 4.4 Derivation of the observed count distributions under the assumption of Beta-distributed capture rates

#### 4.4.1 Telegraph model

If the capture rate *p*_cap_ follows a Beta distribution, *p*_cap_ ∼ Beta(*a, b*), defined by

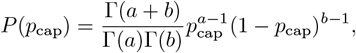

the PGF of the observed counts is given by integrating Eq. (28) over the distribution of *p*_cap_, which yields a generalized hypergeometric function

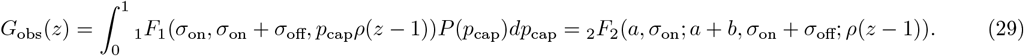

Finally, we obtain the closed-form probability of observing *n* counts

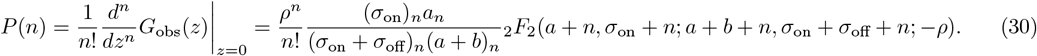

Note that this method of accounting for technical noise, i.e. via compound distributions, is essentially the same as that used to account for static extrinsic noise [98, 99].

#### 4.4.2 Negative binomial model

The PGF of negative binomial distribution, NB(*r, p*), is given by

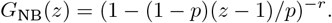

If *p*_cap_ is the same in all cells, the PGF of the observed counts becomes

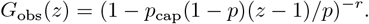

If we have a Beta(*a, b*) distribution of *p*_cap_ then we integrate the above expression over the Beta distribution, yielding

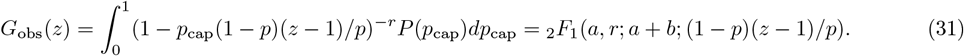

Finally, we obtain the probability distribution of the observed counts

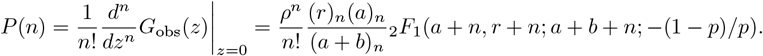

#### 4.4.3 Poisson model

The PGF of the Poisson distribution, Pois(*λ*), is given by

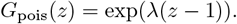

If *p*_cap_ is the same in all cells, the PGF of the observed counts becomes

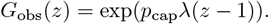

If we have a Beta(*a, b*) distribution of *p*_cap_ then we integrate the above expression over the Beta distribution, yielding

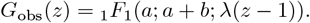

Finally, we obtain the probability distribution of the observed counts

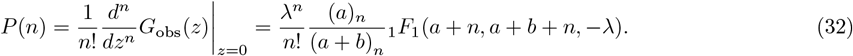

### 4.5 Inference of burst parameters assuming ideal conditions

We consider ideal conditions: (i) infinite sample size so that the measured moments of mRNA numbers do not suffer from finite sample size effects and thus the method of moments can be used for reliable inference; (ii) knowledge of the distribution describing cell-to-cell variability of the capture probability; this means that we can completely correct for technical noise.

#### 4.5.1 Moments of the measured mRNA counts in scRNA-seq data

We assume that the distribution of the measured counts from a gene of interest is given by the telegraph model with capture probability sampled from a Beta(*a, b*) distribution, i.e. Eq. (30).

Given a PGF of observed counts *G*_observed_(*z*), the first two orders of moments can be derived as

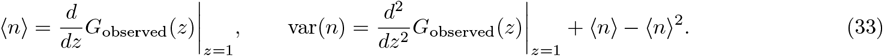

Substituting Eq. (29)into Eq. (33), we obtain expressions for the mean and the variance of the measured count distribution

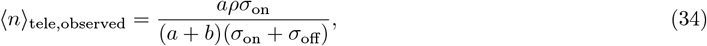

and

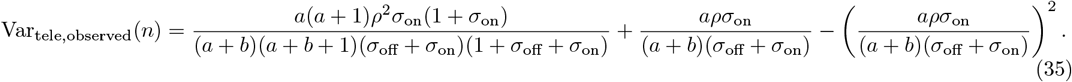

It follows that the Fano factor of mRNA counts (variance divided by the mean) is given by

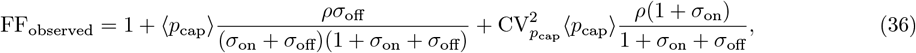

where ⟨*p*_cap_⟩ and 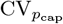 are the mean and the coefficient of variation (standard deviation divided by the mean) of the distribution of the capture probability, Beta(*a, b*).

#### 4.5.2 Moments of the best fit model

We assume that we are in a region of parameter space where aeBIC selects the NB distribution with Beta-distributed capture probability as the optimal model. Substituting Eq. (31) into Eq. (33), we find expressions for the mean and variance of this model

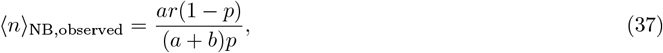

and

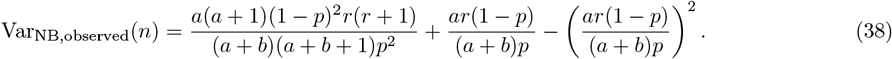

#### 4.5.3 Fitting the model to the data and parameter inference

Since we are assuming that the parameters of the Beta distribution, i.e. *a* and *b*, are known, the only parameters to be inferred are *r* and *p*. These two unknown parameters can be determined exactly by matching the mean and variance of the optimal model Eqs. (37)-(38) with the mean and variance of the data Eqs. (34)-(35). This leads to two simultaneous equations for *r* and *p* which when solved lead to expressions for the inferred two parameters of the NB distribution

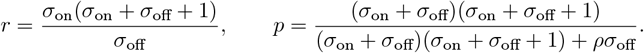

Note that these expressions do not have any dependence on the parameters of the technical noise (*a* and *b*), which indeed show that the inference method has perfectly corrected for technical noise. In fact, *r* and *p* determined in this way are none other than those parameterizing the effective NB distribution Eq. (6) that best fits the conventional telegraph model.

### 4.6 Parameter inference without exact knowledge of the capture rate distribution

Here, we describe an inference approach that simulates the workflow of an actual scRNA-seq experiment including the subsequent analysis of the data. Note that in the inference and model selection parts of the procedure, we do not assume that the distribution of the capture probability is known — rather we use the distribution of the measured total counts per cell as a proxy.

#### 4.6.1 Simulating genomic data

We generated synthetic count data for a genome using the following procedure:

- For each cell *j* = 1, …, *n*_*c*_ (*n*_*c*_ = 1000), sample the capture rate *p*_cap,*j*_ from a Beta(15, 35) distribution which has a mean of ⟨*p*_cap_⟩ = 0.3 and a CV of 0.21. This accounts for cell-to-cell variability in the transcript capture probability.
- Choose the fraction of time spent by genes in the active state from the set *f*_on_ ∈ { 0.1, 0.2, …, 0.7 }, and fix *N*_*σ*_ = 10 and *ρ* = 15. Use the telegraph model as the ground-truth generative model for all genes.
- For each value of *f*_on_ and each cell *j*, sample the transcript counts of gene *i, x*_*ij*_ ∼ Tele(*ρ, f*_on_*N*_*σ*_, (1 − *f*_on_)*N*_*σ*_), independently for 400 genes. This results in transcript numbers per cell for a total of *n*_*g*_ = 2800 genes.
- Simulate observed counts in each cell by downsampling the counts according to the Binomial distribution: *y*_*ij*_ ∼ Binomial(*x*_*ij*_, *p*_cap,*j*_).

#### 4.6.2 Parameter inference and model selection

- Compute the total observed count for each cell *j* as

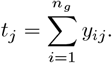
- Estimate the normalization factor *β*_*j*_ using the known mean capture rate ⟨*p*_cap_⟩ (e.g., obtained via spike-in controls) [37]:

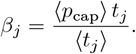
- Use kernel density estimation (KDE) to estimate the distribution *p*(*β*) of the normalization factors.
- For each gene *i*, estimate model parameters by minimizing the negative log likelihood across cells:

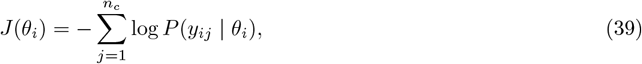

where the likelihood *P* (*y*_*ij*_ | *θ*_*i*_) is computed by marginalizing over *β*:

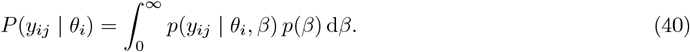

where *p*(*y*_*ij*_ | *θ*_*i*_, *β*) for each candidate model is given by
  – Telegraph: Tele(*ρβ, σ*_on_, *σ*_off_), with *θ*_*i*_ = {*ρ, σ*_on_, *σ*_off_};
  – Negative Binomial: NB(*r*, 1*/*(1 + *bβ*)), with *θ*_*i*_ = {*r, b*}; – Poisson: Pois(*λβ*), with *θ*_*i*_ = {*λ*}.
- Evaluate the integral in Eq. (40) using Gauss quadrature for computational efficiency. Minimize the negative log likelihood using the Nelder-Mead algorithm.
- Compute the Bayesian Information Criterion (BIC) for model selection based on the log-likelihood in Eq. (39).
- If the NB model is selected, estimate the burst size and the burst frequency from the parameter estimates of the telegraph and negative binomial models. For the telegraph model, the burst size is *ρ/σ*_off_ and the burst frequency is *σ*_on_. For the NB model, the estimates are given by 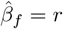 for the burst frequency and 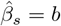 for the burst size.
- Calculate the relative errors in the burst parameter estimates given the actual values used for simulating the genomic data. Randomly select 10^3^ sets of two pairs of burst size and burst frequency estimates. In each set, rank genes by the magnitude of the burst frequency and separately by the burst size. Calculate the percentage of rankings that are flipped compared to the true ranking.

### 4.7 Preprocessing of scRNA-seq dataset

We obtained the raw UMI count matrix for mouse fibroblasts from the file `ss3_n682_fibs_umiCounts.rds`, available at https://github.com/sandberg-lab/lncRNAs_bursting/tree/main/data. To ensure data quality, we applied standard quality control (QC) procedures consisting of two filtering steps. (i) Cell filtering: cells were removed if fewer than 30% of genes had nonzero counts, thereby excluding cells with insufficient transcript coverage. (ii) Gene filtering: genes were removed if they were expressed (nonzero counts) in fewer than 1% of all cells, thereby excluding genes with highly sparse expression. The resulting dataset was then used for downstream statistical modeling and analysis.

### 4.8 MLE-based parameter inference

Given *n*_*c*_ observed mRNA counts 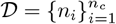, the likelihood of the dataset under kinetic parameters *ϕ* is

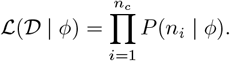

Parameter inference is performed by minimizing the negative log-likelihood,

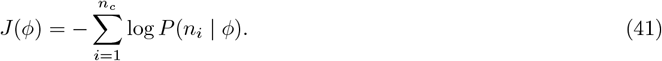

We optimize *J*(*ϕ*) using the Nelder–Mead algorithm implemented in the `Optim.jl` package in Julia. This derivative-free approach avoids costly gradient evaluations of ∂_*ϕ*_*J*(*ϕ*) while maintaining robustness to initialization. Unless otherwise noted, we adopt a gradient tolerance of `g_tol` = 10^−20^ and a maximum of `iterations` = 1000, following the default settings of the `optimize` command.

To account for variability in capture probability *p*_cap_ across cells, we define the normalized capture probability

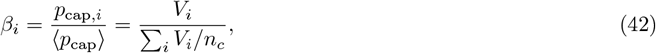

where *V*_*i*_ is the total transcript count of cell *i*. The distribution of *β*, denoted *p*(*β*), is not known *a priori* and is approximated by KDE from the observed values { *β*_*i*_ }, yielding an estimate 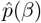.

Efficient computation of *P* (*n*_*i*_ | *ϕ*) requires integrating over the latent variability in *β*. Instead of computing this integral directly, we approximate it using Gauss–Legendre quadrature:

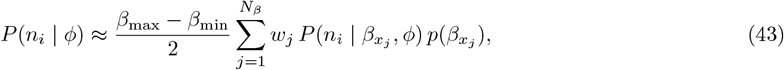

where

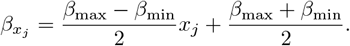

Here *β*_max_ = max_*i*_ *β*_*i*_ and *β*_min_ = min_*i*_ *β*_*i*_, and *x*_*j*_, *w*_*j*_ are the quadrature nodes and weights generated using `gausslegendre`. The probability 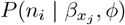 is evaluated using the three *p*_cap_-modified models shown in Fig. 7, with *p*_cap_ replaced by *β*. Under this formulation, the inferred parameters – *ρ* for the telegraph model, *b* for the NB model, and *λ* for the Poisson model – are effectively scaled by the mean capture probability ⟨*p*_cap_⟩, which is typically unknown.

Algorithm 1 summarizes the numerical procedure for MLE under variable capture probability.

#### Algorithm 1

MLE-based parameter inference accounting for *p*_cap_ variability

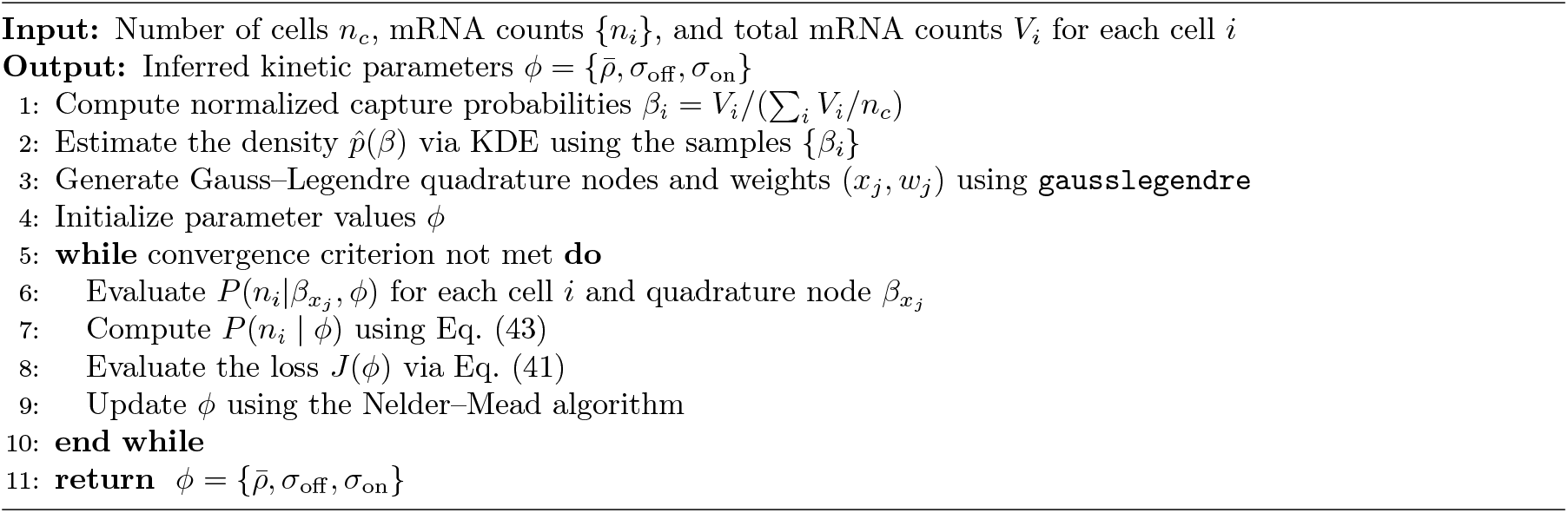

## Supplementary Figures

**Figure S1:**
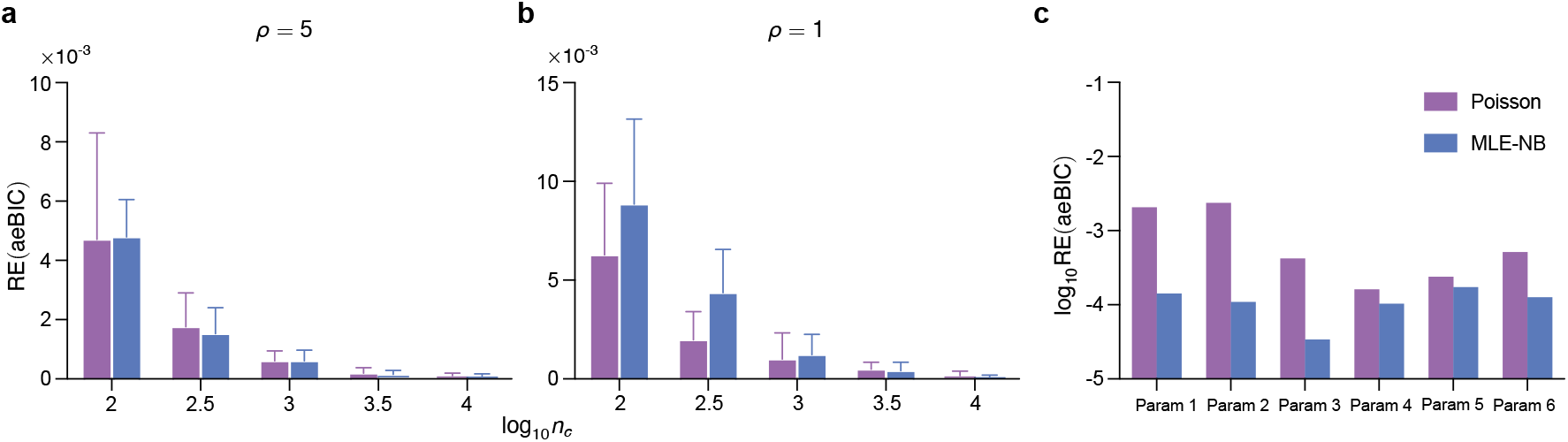
(a)(b) The relative error of aeBIC compared to the expectation of BIC (**E**[BIC]) for two distributions (Poisson and NB) as a function of sample size *n*_*c*_ for 10 parameter sets (see Table 1 for the values of *N*_*σ*_ and *f*_on_) with (a) *ρ* = 5 and (b) *ρ* = 1. Error bars show the standard error of the mean. (c) Relative error of aeBIC with respect to **E**[BIC] for Poisson and NB distributions, evaluated for 6 parameter sets sampled from the plot corresponding to *n*_*c*_ = 10^4^, *ρ* = 15 and *p*_cap_ ∼ Beta(60, 140) in Fig. 6c. The parameter sets are: *f*_on_ = 0.8, *N*_*σ*_ = 3.8 (Param 1), *f*_on_ = 0.88, *N*_*σ*_ = 1.5 (Param 2), *f*_on_ = 0.5, *N*_*σ*_ = 38 (Param 3), *f*_on_ = 0.96, *N*_*σ*_ = 758.6 (Param 4), *f*_on_ = 0.7, *N*_*σ*_ = 380.2 (Param 5), *f*_on_ = 0.1, *N*_*σ*_ = 631 (Param 6).

**Figure S2:**
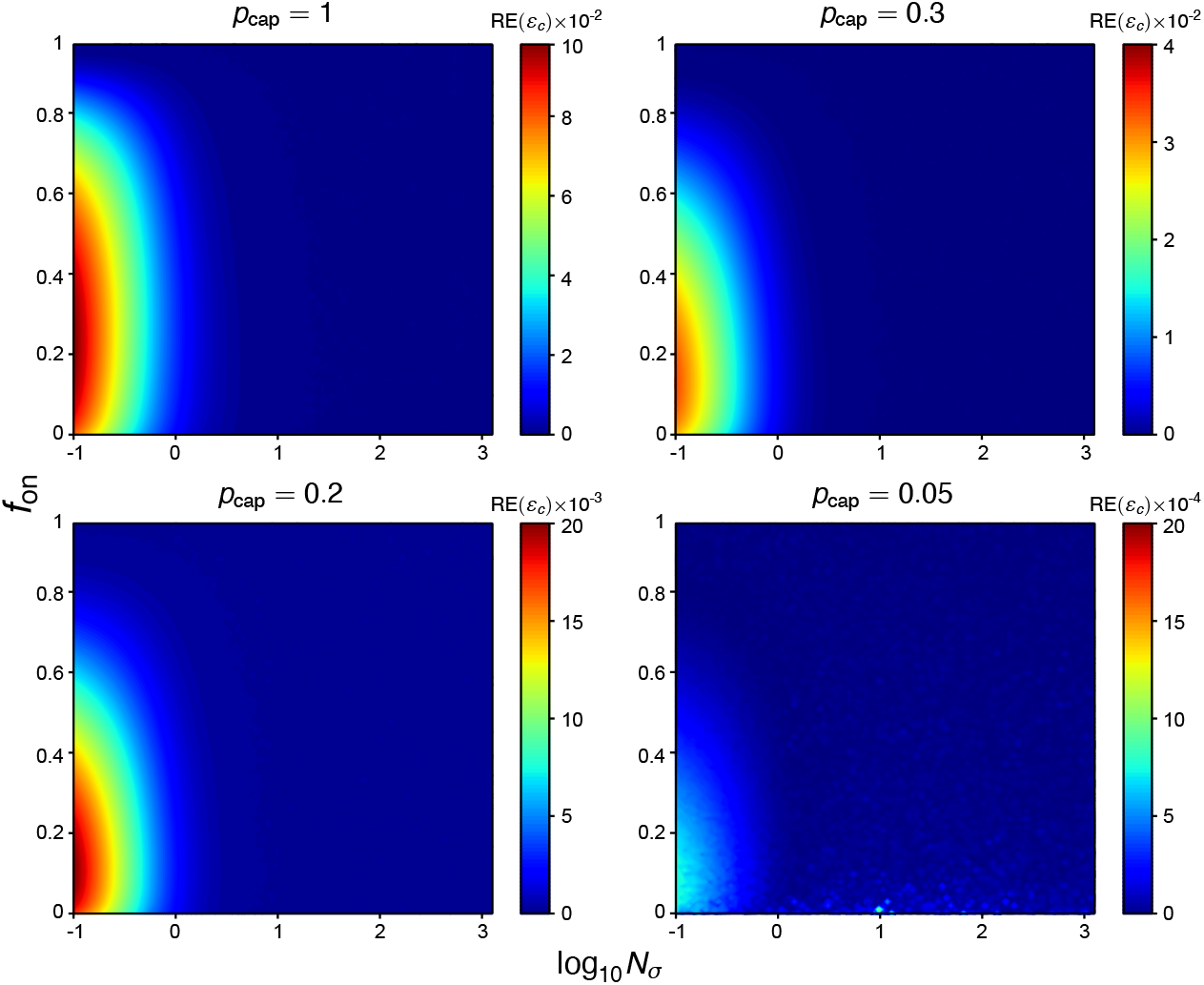
The relative error between the minimum cross-entropy determined by MLE and moment matching. The cross-entropy is that between the NB distribution as the proposed model (*P*_ℳ_ (*n* | *θ*)) and the telegraph model distribution Eq. (1) with *ρ* → *ρp*_cap_ as the ground-truth distribution (*P*^𝒢^ (*n*)), assuming all cells have an identical mRNA capture probability *p*_cap_. The minimum cross-entropy using MLE is obtained by finding the two parameters of the NB distribution which minimize Eq. (11). The minimum cross-entropy using moment-matching is obtained by directly evaluating Eq. (11) using the effective NB distribution (Eq. (6)) with *ρ* → *ρp*_cap_. As the heat maps show, the relative error between the two minimum cross-entropies is very small, regardless of the values of *p*_cap_, *f*_on_ and *N*_*σ*_. Note that *ρ* is in all cases fixed to 15.

**Figure S3:**
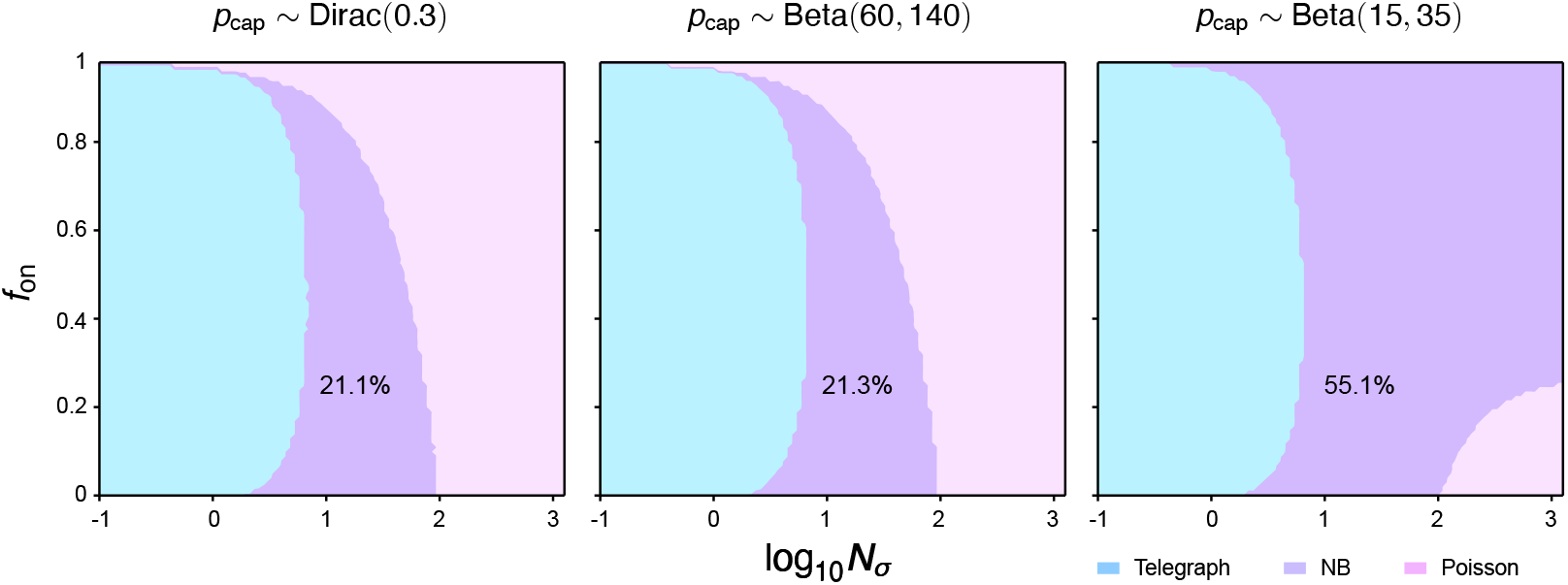
Phase diagram showing the regions of parameter space where the technical-noise corrected telegraph, NB and Poisson models are selected as the optimal models by the aeBIC, given that the ground-truth mRNA distribution is that of the corrected telegraph model, i.e. the telegraph model with *p*_cap_ sampled from Dirac(0.3), Beta(60, 140) and Beta(15, 35) — these distributions have mean 0.3 and CV = 0, 0.11 and 0.21, respectively. Here *N*_*σ*_ is the sum of gene-state switching rates normalised by the degradation rate of mRNA, and *f*_on_ is the fraction time spent in the active state. The fraction of the total parameter space occupied by the region where the corrected NB distribution is optimally selected is shown on the plots. Note that the transcription rate is fixed to *ρ* = 15. The maximum mean number of transcripts in the phase plots is 4.5. The sample size is fixed to *n*_*c*_ = 10^4^.

**Figure S4:**
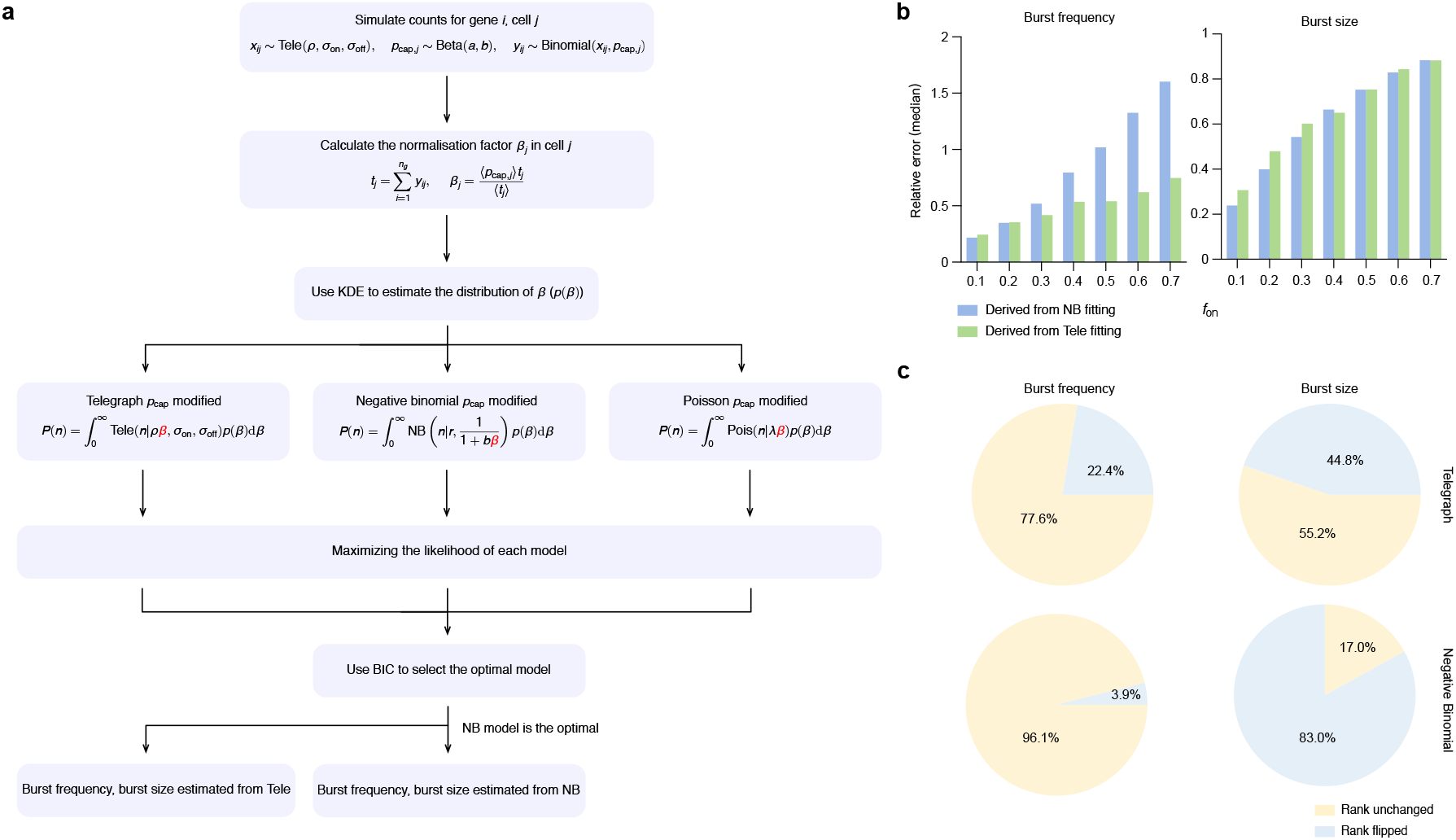
(a) Flowchart of the procedure described in Section 4.6 which simulates the workflow of a scRNA-seq experiment. A synthetic scRNA-seq dataset (1000 cells and 2800 genes) is simulated — technical noise is account for by sampling the capture probability for cell *j* (*p*_cap,*j*_) from a Beta(15, 35) distribution. The data is then used to perform MLE-based parameter inference and BIC-based model selection. Notably, the latter is done without exact knowledge of the capture rate distribution, which mimics the real scenario. Only knowledge of the mean capture probability *p*_cap_ is assumed which can be estimated using spike-in controls [8] or by comparison of scRNA-seq data with smFISH data [10]. (b) Plots of the median relative error of the estimates of the burst frequency and burst size as a function of the true fraction of time spent in the active state *f*_on_ by genes. The estimates are for those genes whose distribution of counts per cell is best fit by an NB distribution (after correcting for technical noise). The estimates are obtained by maximizing the likelihood of the technical-noise corrected NB and telegraph models (blue and green bars, respectively). (c) 10^3^ sets of two pairs of burst size and burst frequency estimates are randomly selected. In each set, genes are ranked by the magnitude of the burst frequency and separately by the magnitude of the burst size. The piecharts show the percentage of rankings that are flipped compared to the true ranking. The rankings are separately done using the parameter estimates obtained by maximizing the likelihood of the technical-noise corrected NB and telegraph models (piecharts in the bottom and top rows, respectively).

**Figure S5:**
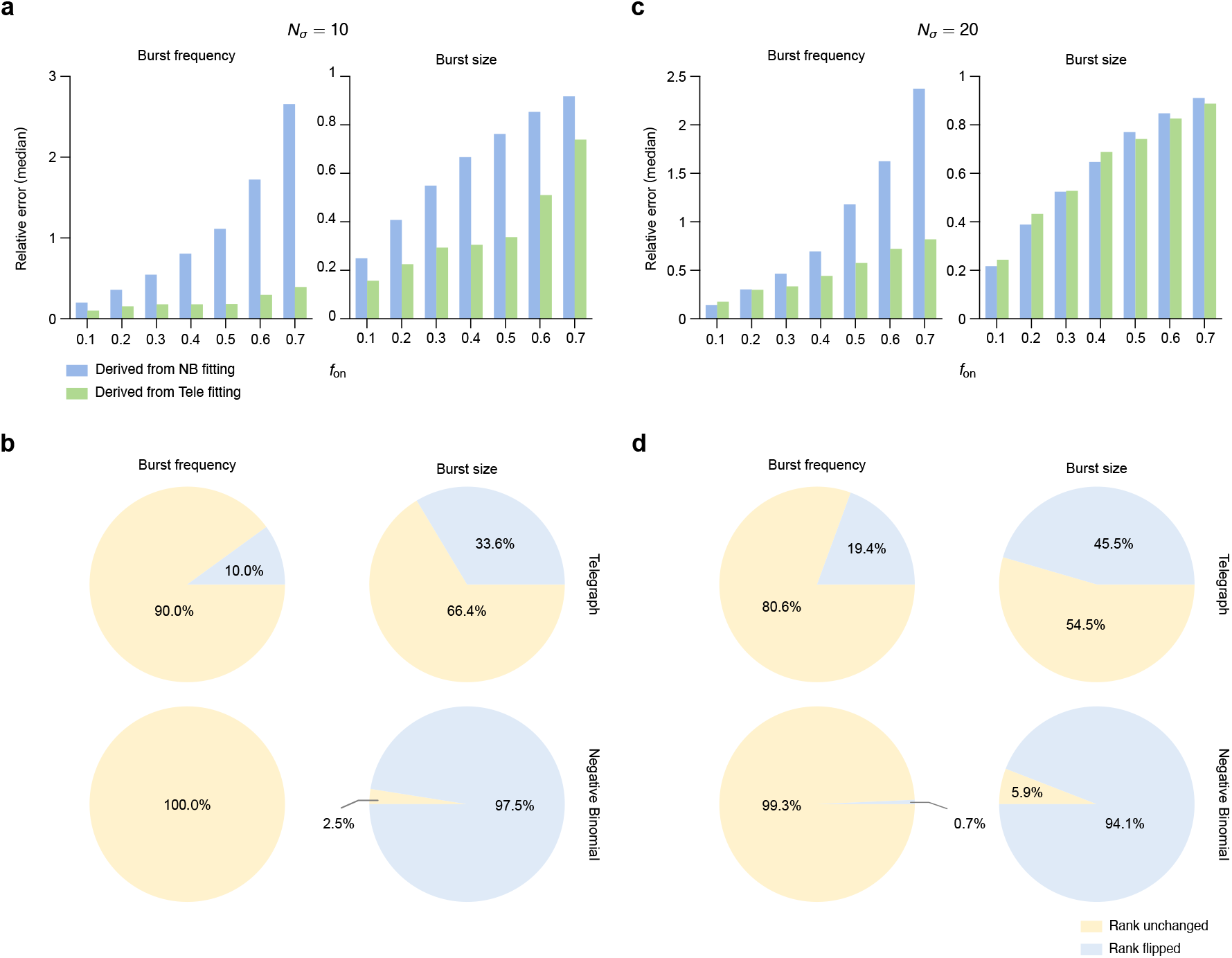
(a)(b)Recalculation of the statistics shown in Fig. S4(b)(c) using the procedure described in Section 4.6 with the difference that the number of cells is *n*_*c*_ = 10^4^ and the transcript capture distribution is sampled from Beta(60, 140). Note that *N*_*σ*_ = 10. (b)(d) Recalculation of the statistics shown in (a) and (b) with the difference that *N*_*σ*_ = 20. Note that 10^3^ sets of two pairs of burst size and burst frequency estimates were used to compute the piecharts for the gene rankings in (b) and (d).

**Figure S6:**
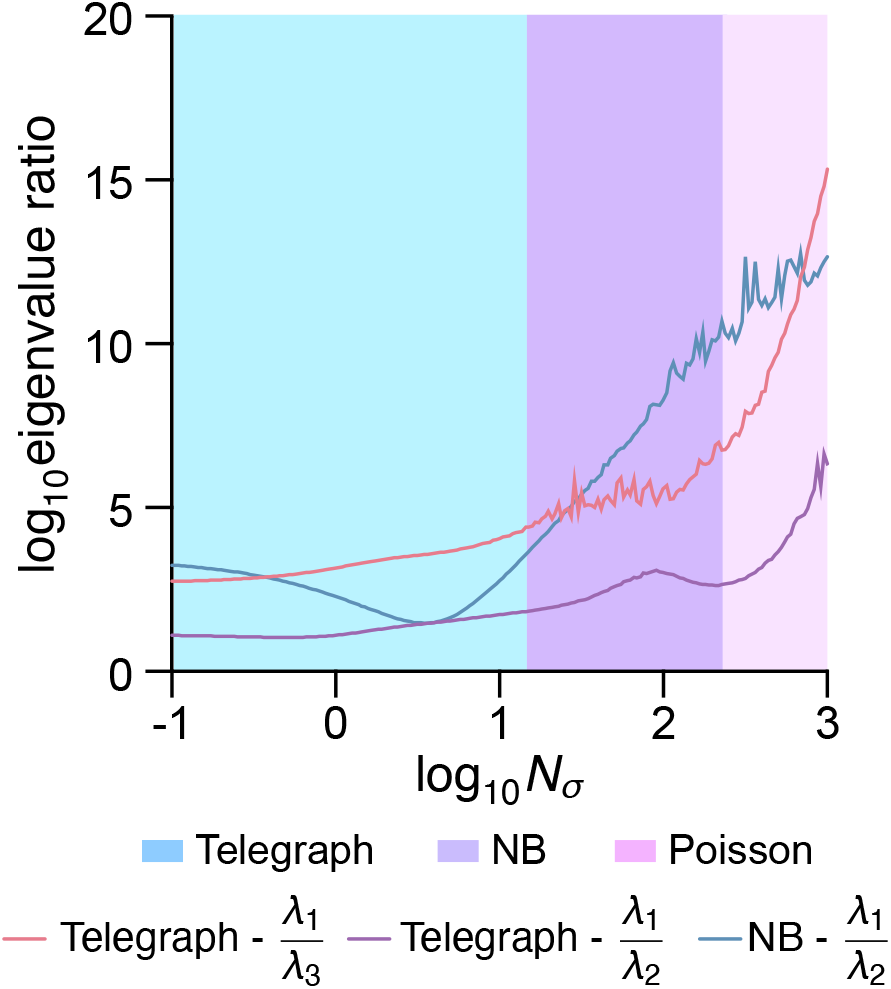
Eigenvalues of the Hessian matrix of the log-likelihood are not reliable indicators for model selection. We varied *N*_*σ*_ from 0.1 to 10^3^, while fixing *ρ* = 15, *f*_on_ = 0.3, and the number of cells at 10^4^. The corresponding values of *σ*_on_ and *σ*_off_ were determined by Eq. (7) in the main text. For fair comparison with the aeBIC method, data were generated directly from Eq. (1) using these parameters. The log-likelihood was defined as ℒ_*θ*_ = *n*_*c*_ ∑_*n*_ *P* (*n*) ln *P*_*θ*_(*n*), where *P*_*θ*_(*n*) is the probability given by either the telegraph or NB model with parameters *θ*. Under this setup, the telegraph model recovers the generative parameters exactly, while the NB parameters follow Eq. (6). We then computed the eigenvalues of the Hessian matrix at the MLEs for each model: three eigenvalues for the telegraph model and two for the NB model, all negative. Denoting the ordered eigenvalues as *λ*_1_, *λ*_2_, *λ*_3_, we examined the ratios − *λ*_1_*/λ*_2_ and − *λ*_1_*/λ*_3_ for the telegraph model, and − *λ*_1_*/λ*_2_ for the NB model, as *N*_*σ*_ increased. These ratios (lines) gradually become infinite as *N*_*σ*_ grows, reflecting model degeneration. In contrast, the aeBIC method yields clear model selection boundaries (shaded areas). Together, these results indicate that Hessian eigenvalues are not reliable indicators for model selection.

